# The Unique Efg1 Fungal Virulence Regulon in the Catheterized Bladder Environment

**DOI:** 10.1101/2025.11.06.686417

**Authors:** Alyssa Ann La Bella, Nicholas C. Gervais, Kurt N. Kohler, Chloe L.P. Obernuefemann, Ilse D. Jacobsen, Rebecca S. Shapiro, Michael G. Caparon, Scott J. Hultgren, Ana Lidia Flores-Mireles, Felipe Hiram Santiago-Tirado

## Abstract

<u>Urinary</u> catheterization, a common procedure in hospitals and nursing home facilities, is a primary driver of hospital-acquired infections (HAI). These devices frequently lead to catheter-associated urinary tract infections (CAUTIs), which often progress to severe complication, sepsis, and ultimately death. The fungus *Candida albicans* has emerged as the second most common causative agent of CAUTIs; yet, its pathogenesis is poorly understood, which complicates development of efficient treatments. Previously, we identified the transcription factor Efg1 as a critical virulence driver in *C. albicans* CAUTIs. However, its specific downstream targets within the unique bladder microenvironment remained unknown. This study identifies, for the first time, the complete Efg1 regulon that is active during growth in human urine. We confirmed the clinical relevance of this discovery, finding that many of these Efg1-regulated factors are present and significantly upregulated in catheter samples from patients with *C. albicans* infections. Furthermore, we characterized two of these key factors, *ECE1* and *EED1*, validating their roles both *in vitro* in urine conditions and *in vivo* using a CAUTI mouse model. Identifying the tissue-specific downstream targets of Efg1 elucidates the precise mechanism of fungal CAUTI. This knowledge provides a new roadmap for developing targeted therapeutics, offering vital antimicrobial-sparing strategies to combat these life-threatening infections.

**SIGNIFICANCE:** Catheter-associated urinary tract infections (CAUTIs) are common hospital-acquired infections that can lead to severe complications and death. Although most are caused by bacteria, the fungus *Candida albicans* is an increasingly prevalent cause, yet the pathogenesis of fungal CAUTIs is poorly understood. Previous research identified Efg1 as necessary for CAUTI, and now this study defines the urine-specific Efg1 regulon, validating its clinical relevance in catheter samples from infected patients. We further assessed how key downstream factors, Ece1 and Eed1, contribute to bladder infection. This first report of the urine-specific *EFG1* network provides new targets for diagnosing and treating these life-threatening infections.

## INTRODUCTION

In 2022, the World Health Organization classified *Cryptococcus neoformans*, *Candida auris*, *Aspergillus fumigatus*, and *Candida albicans* as critical fungal threats, focusing global attention on these pathogens (1). This “critical priority” status stems from their significant medical burden, increasing antifungal resistance, and the limited availability of effective therapies (1). Among these, *C. albicans* is the most common causative agent of invasive fungal infection. However, the true global burden of these diseases remains poorly defined, as fungal pathogens have historically received insufficient resources and scientific consideration (1).

*C. albicans’* pathogenicity is driven in part by its ability to form biofilms (2, 3), which it readily establishes on indwelling medical devices like urinary catheters, pacemakers, mechanical heart valves, joint prostheses, contact lenses, and dentures (1–3). These biofilms pose a significant clinical challenge as they are notoriously resistant to antifungal treatments and capable of evading the host immune response. This resilience is a primary contributor to persistent fungal infections and the growing threat of antifungal resistance (4).

Between 15 and 25% of hospitalized patients receive a urinary catheter, predisposing them to catheter-associated urinary tract infections (CAUTIs), one of the most common hospital-acquired infections (HAI) (5–9). While most CAUTIs are bacterial, *Candida* species (spp.), particularly *C. albicans*, have become a primary causative agent, second only to uropathogenic *E. coli* in prevalence (7). CAUTI pathogenesis is known to rely on the human wound-healing protein, fibrinogen (Fg), which serves as a scaffold for biofilm formation and a nutritional source for numerous pathogens in the catheterized bladder (9–11). *C. albicans* CAUTIs require both hyphal formation, which is induced by the urine environment, and the ability to adhere to Fg, which promotes biofilm formation (11). These bladder infections are a significant threat because they can disseminate systemically into the bloodstream, leading to potentially fatal outcomes (12, 13). As ∼25% of sepsis cases in hospital and nursing home settings begin in the urinary tract (urosepsis), understanding the mechanisms of these bladder-initiated infections is a critical public health priority (14, 15). Thus, we have begun to dissect the pathophysiology of fungal CAUTIs (11, 16).

Previously, we found that the transcription factor Efg1 (Enhanced filamentous growth 1) is responsible for hyphal morphogenesis in the catheterized bladder environment (11). Furthermore, we identified the Agglutinin-like sequence 1 (Als1) as the main adhesin for Fg in urine microenvironment and, importantly, as a downstream effector of *EFG1* (11, 17). In a CAUTI mouse model, deletions of either *EFG1* or *ALS1* resulted in defective colonization(11). The *als1*Δ*/*Δ strain, which could still form filaments, was unable to colonize the catheterized bladder, confirming that adhesion is crucial in the dynamic and open bladder environment (11). To test if Als1 was sufficient to rescue the *efg1*Δ*/*Δ defect, we overexpressed *ALS1* in the *efg1*Δ*/*Δ strain (*efg1*Δ*/*Δ + pALS1) in our urine-Fg dependent biofilm formation and our mouse model of CAUTI (11). We found that Als1 overexpression was unable to fully restore biofilm formation or bladder and catheter colonization to wild-type (WT) strain (11). This indicates that in addition to Efg1-dependent hyphal morphology and Als1-Fg-dependent adhesion, other Efg1-regulated factors are critical for *C. albicans* persistence during fungal CAUTI.

Efg1 is a critical *C. albicans* transcription factor involved in numerous processes including morphogenesis, white-opaque switching, metabolism, and biofilm formation (18–21). Efg1 targets are known to be condition-dependent, and its regulation can differ between clinical isolates (22, 23). While Efg1 has been studied in gut and oropharyngeal infections, its regulatory network within the urinary tract (or urine) environment has remained unexplored (24, 25).

In this study, we identify the Efg1-regulated factors in the unique urine microenvironment. Importantly, we validated key factors in *efg1*Δ*/*Δ Fg-urine biofilms and confirmed their clinical relevance, finding them upregulated in *C. albicans*-infected patient catheters during mono- and polymicrobial infections. We further characterized *ECE1*, which encodes the epithelial-damaging toxin candidalysin, and *EED1* (also known as *DEF1*), which is essential for hyphal extension. The role of these genes was investigated *in vitro* and *in vivo* using deletion and overexpression mutants, finding that although dispensable for CAUTI establishment, they play distinct roles. Understanding these Efg1-regulated factors is crucial for elucidating the mechanisms of these ever-increasing nosocomial fungal infections.

## RESULTS

### Identification of *C. albicans* deletion mutants with significantly enhanced or defective Fg-urine dependent biofilms

Using our well-established Fg-urine biofilm assay, we screened three *C. albicans* homozygous deletion mutant libraries made in the SC5314 strain background: **1)** Suzanne Noble’s collection (26); **2)** Oliver Homann’s transcription factor (TF) deletion library (27); and **3)** Rebecca Shapiro’s adhesin deletion library (28). Of the 687 unique mutants, we found that 155 form biofilms significantly different from WT: 142 mutants (red) exhibited a defect in biofilm formation, and 13 mutants (green) increased biofilm formation (**Fig. 1A**). Of all the mutants, *efg1*Δ*/*Δ and *als1*Δ*/*Δ showed the most severe Fg-urine biofilm defect, validating our previous findings (11). We performed STRING network analysis on the genes involved in either promoting (**Fig. 1B, Table S1**) or inhibiting (**Fig. 1C, Table S2**) biofilm formation under Fg-urine conditions. Seven of the 13 factors that promoted biofilm formation had previously no known interactions (**Fig. 1B**) and 53 of the 142 mutants that resulted in defective biofilm formation had previously no known interactions (**Fig. 1C**). However, the network of deficient biofilm formers had an overall higher number of edges (151), or protein interactions, than expected (50), indicating that these factors may be partially biologically connected in the urine environment (**Fig. 1C, Table S2**). Furthermore, *EFG1* has many direct and indirect interactions with factors that resulted in defective biofilm formation when deleted (**Fig. 1C, Table S2**). Gene ontology (GO) analysis of the biological processes of the 155 significant factors showed that they have diverse roles in interspecies interactions, stress response, filamentation, cell wall organization, and adhesion (**Fig. 1C and D, Table S3**).

**Figure 1.**
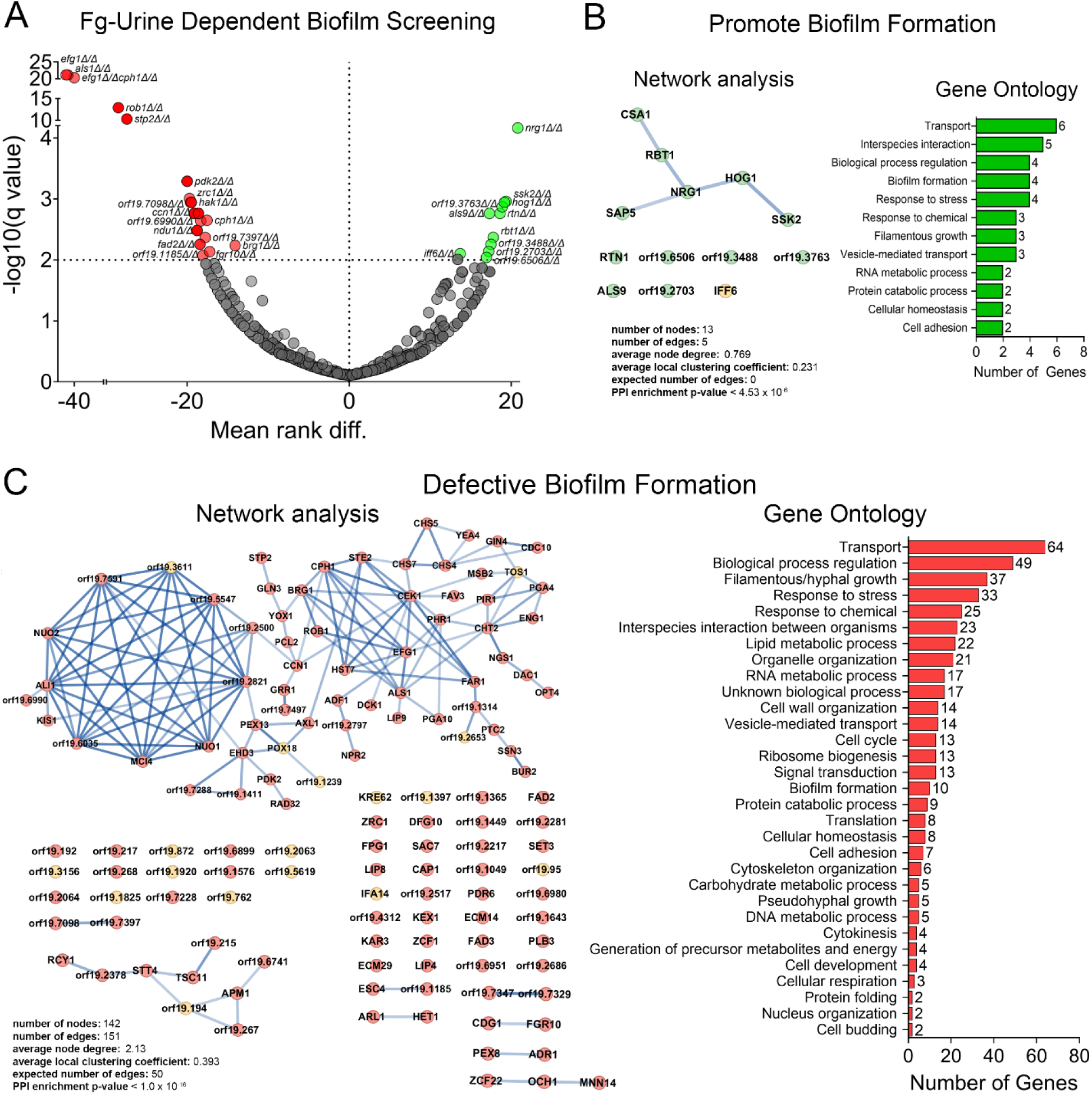
*C. albicans* deletion mutants in the Fg-urine biofilm model show significant defective (142 mutants) and enhanced (13 mutants) biofilm formation phenotypes. (**A**) Fg-urine biofilm formation screening of deletion mutants reveals 155 statistically significant mutants that either enhanced (13 mutants, green) or resulted in defective (142 mutants, red) biofilm formation compared to WT. (**B**) STRING network analysis of the deletion mutants that promoted Fg-urine biofilm formation. Circles represent nodes, or proteins. Lines represent edges and indicate interaction between proteins. Line darkness indicates confidence score or strength of interaction. Yellow nodes indicate uncharacterized proteins. Biological process gene ontology analysis of the factors that had a statistically significant increase in biofilm formation. (**C**) STRING network analysis of the deletion mutants that resulted in defective Fg-urine biofilm formation. Circles represent nodes, or proteins. Lines represent edges and indicate interaction between proteins. Line darkness indicates confidence score or strength of interaction. Yellow nodes indicate uncharacterized proteins. Biological process gene ontology analysis of the factors that had a statistically significant decrease in biofilm formation.

### The urine environment induces a unique Efg1 regulon

Based on the severe biofilm defect of *EFG1* deletion mutant and that Efg1 is a transcriptional regulator linked to many biofilm formation factors (**Fig. 1**), we prioritized defining the *EFG1* regulon in urine conditions to elucidate the mechanisms of *C. albicans* CAUTIs. To unveil this regulon, we extracted total RNA from WT, *efg1*Δ*/*Δ, or *EFG1*-complement (*efg1*Δ/Δ+*EFG1*) strains grown in Fg-dependent biofilms in urine or YPD media for 48 hours. RNA samples were subject to RNA sequencing, and the transcriptional profile of *C. albicans efg1*Δ*/*Δ grown in Fg-urine-dependent biofilms was compared to the transcriptional profile of WT Fg-urine biofilms (**Fig. 2A, Table S4**). The data from the *EFG1*-complement strain (**Fig. S1, Table S5 and S6**) was used to ensure that any dysregulation is not due to polar effects caused by genetic manipulation of the *efg1*Δ*/*Δ strain (**Fig. S1**, **Table S5 and S6**). The genes found to be exclusively downregulated in *efg1*Δ*/*Δ Fg-urine biofilm RNA were thus determined to be a part of the Efg1-urine regulon.

**Figure 2.**
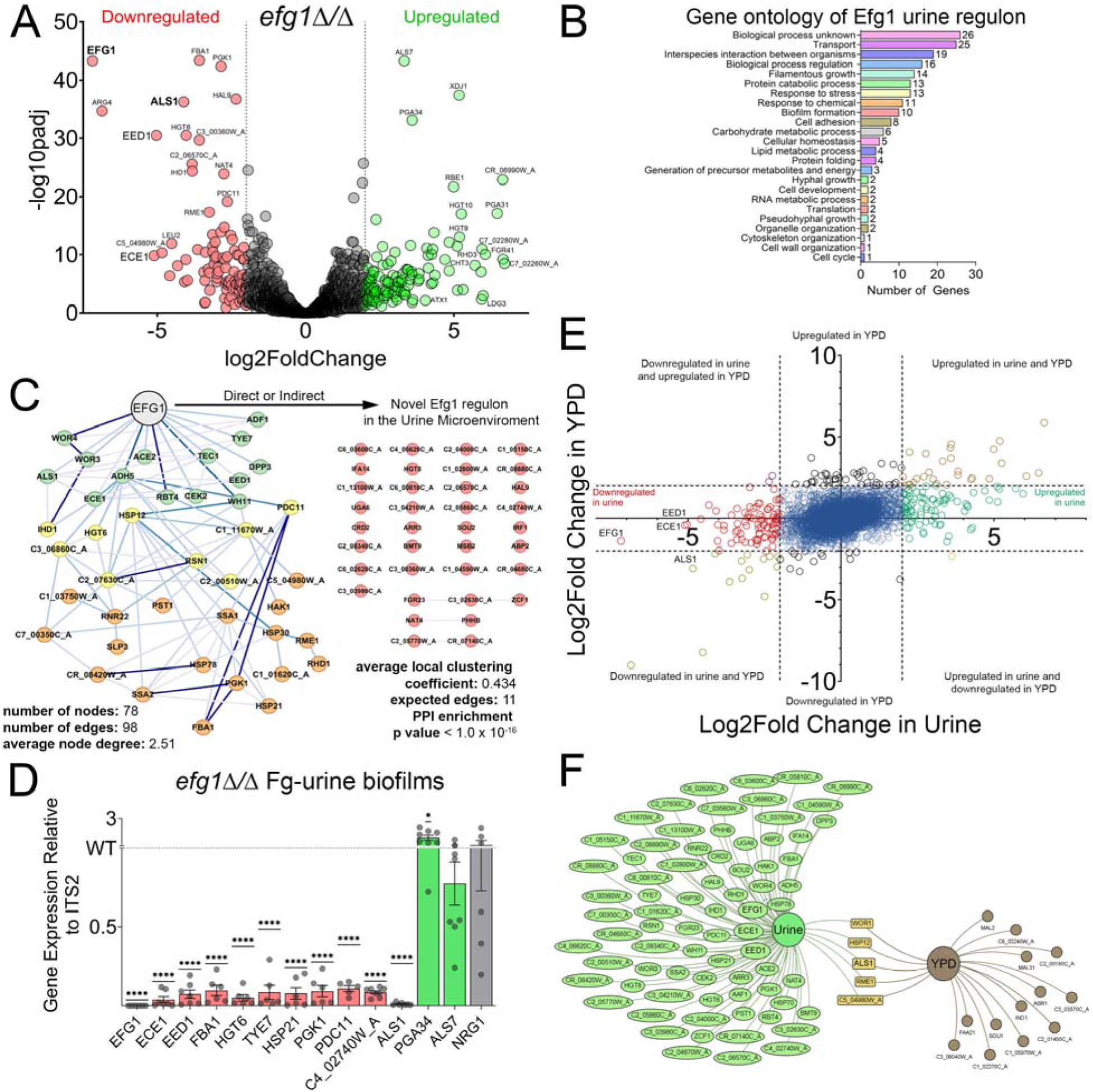
The Efg1 Regulon in the Urine Microenvironment. (**A**) Volcano plot showing downregulated (red, log2FoldChange ≤ −2) and upregulated (green, log2FoldChange ≥ 2) factors in Fg-dependent urine *efg1*Δ*/*Δ biofilms from RNA sequencing. (**B**) Biological processes gene ontology analysis of genes downregulated in Fg-dependent urine *efg1*Δ*/*Δ biofilms (Efg1-regulated genes in urine). (**C**) STRING network analysis showing protein-protein interactions of the Efg1 urine-specific regulon. Circles represent nodes, or proteins. Lines represent edges and indicate interaction between proteins. Line darkness indicates confidence score or strength of interaction. (**D**) Validation of Efg1-urine regulon in Fg-urine dependent biofilms via qRT-PCR. RNA was extracted from SC5314 WT and *efg1*Δ*/*Δ 48-hour Fg-urine biofilms, and expression of select Efg1-urine regulon genes were found by qRT-PCR and analyzed with the 2^-ΔΔCt^ method. Data are shown as the relative expression normalized to the housekeeping gene, *ITS2*, and SC5314 WT. (**E**) Scatter plot comparing the log2 fold change of all genes between efg1Δ*/*Δ and WT under urine (X-axis) and under YPD (Y-axis) conditions. In red are the genes uniquely downregulated in *efg1*Δ*/*Δ under Fg-urine conditions, and the ones downregulated under both conditions are shown in yellow (most of which are the auxotrophic markers in SN250). (**F**) Venn diagram showing the number of genes downregulated in *efg1*Δ*/*Δ compared to WT under urine or YPD conditions. Stringent thresholds of at least 4-fold change and no change in the *efg1*Δ*/*Δ complemented strain were used to identify differentially regulated genes (see Methods). Only 5 genes were downregulated in *efg1*Δ*/*Δ under both conditions when comparing urine to YPD, allowing us to identify the urine-Fg specific Efg1 regulon.

Several notable genes were found, including *ALS1* (the main Fg adhesin in urine conditions), *EED1* (involved in filament elongation and hyphal maintenance), *ECE1* (encoding the candidalysin toxin), and *WOR3* (required for white-opaque switching/mating) (11, 20, 29, 30) (**Fig. 2A, Table S4**). GO analysis of biological processes revealed that 26 genes were uncharacterized, indicating that many factors relevant to fungal CAUTIs remain unknown (**Fig. 2B, Table S9**). Other genes were classified as critical for host interactions, stress response, filamentous growth, and biofilm formation (**Fig. 2B**). Furthermore, a STRING protein-protein association network analysis confirmed many previously known gene interactions with *EFG1* (**Fig. 2C, Table S10**), including a direct association with *ALS1*, which validates our previous findings (11). Importantly, we also discovered novel associations that were not previously known for Efg1, of which 26 are uncharacterized genes (**Fig. 2C, Table S9**).

We validated our RNA-seq findings via qRT-PCR by selecting ten genes (*ECE1*, *EED1*, *FBA1*, *HGT6*, *TYE7*, *HSP21*, *PGK1*, *PDC11*, *C4_02740W_A, ALS1*; red) based on their wide-ranging roles in virulence, carbohydrate metabolism, filamentation, biofilm formation, adhesion, and transport. We confirmed their downregulation in *efg1*Δ*/*Δ Fg-urine biofilms, confirming Efg1-dependent regulation (**Fig. 2D**). We also validated two upregulated genes (*PGA34*, *ALS7*; green) (**Fig. 2D**). Notably, deletion of *NRG1*, an Efg1 repressor, promotes Fg-urine biofilm formation (**Fig. 1A**). Since other have shown that *nrg1*Δ*/*Δ increases hyphal formation (11, 31), we tested its expression under *efg1*Δ*/*Δ Fg-urine conditions, finding that *NRG1*’s expression was similar to WT levels (**Fig. 2D**). This highlights the importance of other transcriptional regulators in the catheterized bladder. Our transcriptomic findings were further confirmed in an independently-created CRISPR *efg1*Δ*/*Δ mutant (SC5314 background – see Methods for more information) (**Fig. S3**).

Of the Efg1-urine regulon factors, we aimed to identify the factors that were unique to urine, as opposed to also being Efg1-regulated in Fg-YPD conditions. To identify Efg1 factors regulated uniquely in urine, the downregulated genes in *efg1*Δ*/*Δ urine biofilms relative to WT-urine and the downregulated genes in *efg1*Δ*/*Δ-YPD biofilms relative to WT-YPD were compared (**Table S4-7**). Eighty factors were downregulated in *efg1*Δ/Δ-urine when compared to WT-urine, and only 5 of those were also downregulated in YPD conditions. This is not surprising given the limited role Efg1 plays under YPD conditions. Thus, we identified 75 factors uniquely Efg1-regulated in Fg-urine conditions (**Fig. 2E-F, Table S8**).

### Clinical catheter isolates show upregulation of factors in the Efg1 urine regulon

Building on our animal model, which accurately mimics human infection (9, 10, 32–37), we next sought to determine the role of Efg1 in clinical CAUTI. For this, we analyzed *C. albicans* transcriptional profiles from CAUTI patient’s catheters via RNA sequencing. The patient cohort was diverse, including both male and female patients who had varying health conditions, catheter dwell times, and microbial co-occurrences (**Fig. 3A**). HUC123-02, HUC123-03, and HUC123-04 were longitudinal catheters from one patient. Notably, only catheter HUC103 did not have a co-occurrence with another microbe (**Fig. 3A**). Hierarchical clustering showed that fungal transcriptomes were highly similar, with the exception of the monomicrobial HUC103 (**Fig. 3B**). We then focused on the Efg1-urine regulon (from **Fig. 2**) within the fungal transcriptomes from the catheterized patients. Non-metric multidimensional scaling (MDS) using the Bray-Curtis coefficient revealed ∼80% similarity among these regulon profiles (**Fig. 3C**). Furthermore, the fungal transcriptional profiles from the three catheters from patient HUC123 showed ∼90% similarity to one another (**Fig. 3C**). This data suggests that Efg1 and its downstream targets in urine remain relevant in clinical settings.

**Figure 3.**
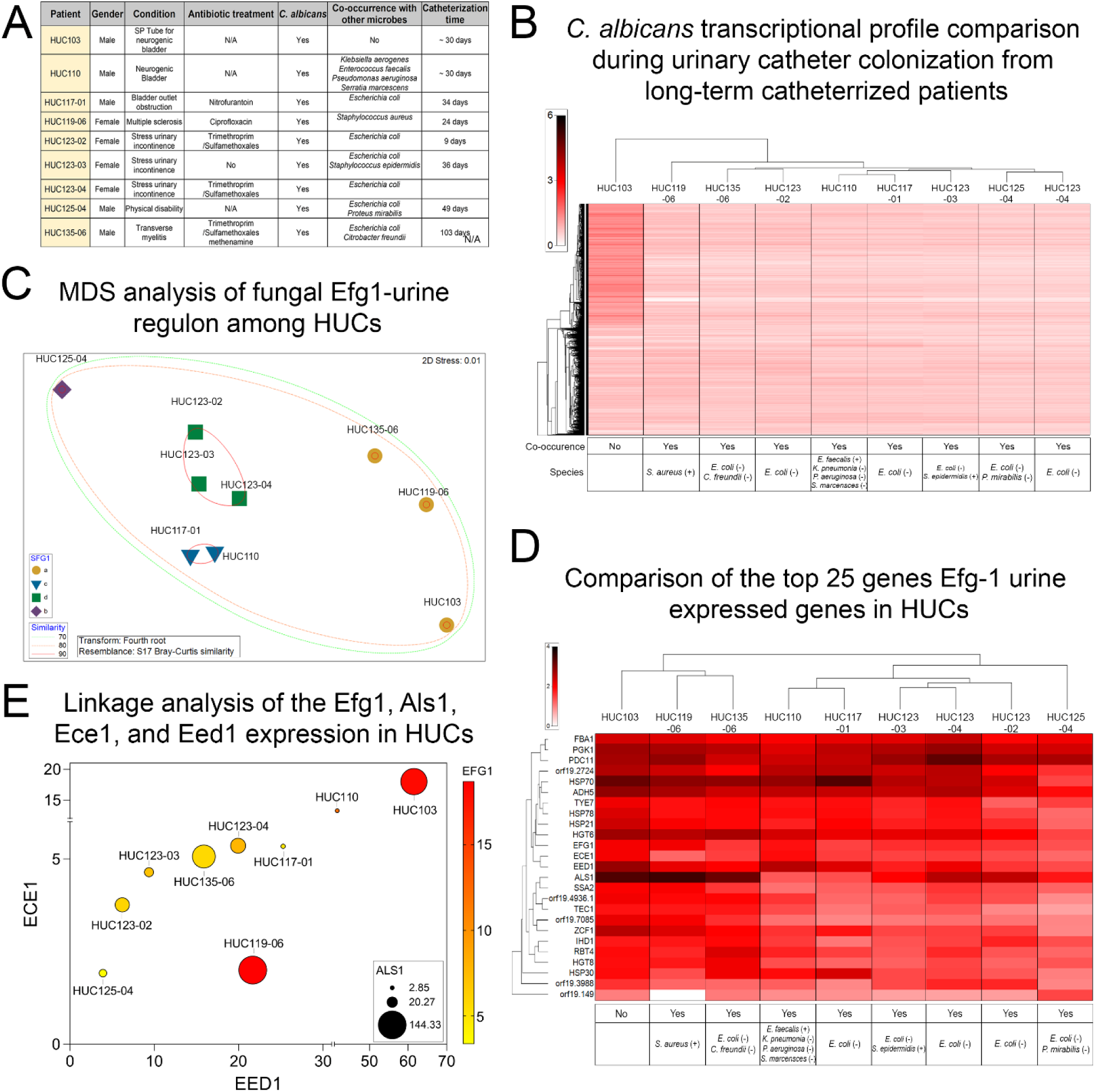
The Efg1-urine regulon is upregulated in RNA isolated from patient catheters with *C. albicans* infection. (**A**) Patient data for urinary catheters collected and analyzed in this study. (**B**) Comparison of the expression profiles of the entire genomes of the clinical urinary catheters along with their co-isolated pathogens. (**C**) Multivariable data analysis comparing the expression of Efg1-urine specific downstream factors across clinical catheter samples. Circle sizes correspond with relative Efg1 expression. Dotted colored lines indicate the level of similarity between enclosed isolates (70%, green; 80%, orange; 90%, red). (**D**) Expression profile of the top 25 Efg1-urine regulon factors from clinical urinary catheters along with their co-isolated pathogens. (**E)** Multivariable analysis showing the clinical catheter isolates’ *EFG1*, *ALS1*, *ECE1*, and *EED1* expression. Color corresponds to *EFG1* expression, circle size corresponds to *ALS1* expression, x-axis corresponds to *EED1* expression, and y-axis corresponds to *ECE1* expression.

Additionally, the expression of the top 25 Efg1-urine regulated genes for each catheter was compared via cluster analysis, revealing two main groups (**Fig. 3D**). The first cluster grouped HUC103 (*C. albicans* only) with HUC119-06 and HUC135-06 (co-occurrences with *Staphylococcus aureus, Escherichia coli,* and *Citrobacter freundii*). The second cluster contained HUC110, HUC117-01, HUC123-03, HUC123-02, HUC123-04, and HUC 125-04 (co-occurrences with *E. coli, Klebsiella pneumonia, Pseudomonas aeruginosa, Serratia marcensces, S. epidermis,* and *Proteus mirabilis*) (**Fig. 3D**). These group separations suggest that co-occurrence with other microbes may modulate *C. albicans’* Efg1-urine regulon and its pathogenesis.

The Efg1-urine regulated factors, *ECE1* (<u>e</u>xtent of <u>c</u>ell <u>e</u>longation protein 1; encoding the candidalysin toxin) and *EED1* (<u>e</u>pithelial <u>e</u>scape and <u>d</u>issemination protein 1; needed for maintenance of hyphal growth) have been characterized in other model systems (29, 30, 38–41), but their role during fungal CAUTIs is unknown. To begin understanding their contribution to infection, we performed a correlation analysis of the relative expression of *EFG1*, *ALS1, ECE1,* and *EED1* for each human catheter (**Fig. 3E**). Overall, the expression of these genes was directly correlated. Interestingly, the monomicrobial HUC103 sample had the greatest overall expression of *EFG1*, *ALS1*, *ECE1*, and *EED1* (**Fig. 3E**). The other polymicrobial catheters had lower expression of these genes compared to HUC103, although their expression was still upregulated. This finding further suggests that the presence of other microbes modulates the transcriptional profile of *C. albicans* during CAUTI (**Fig. 3D and E**).

### Ece1 and Eed1 mutants show phenotypic variation in urine *in vitro* conditions

Our RNA sequencing and qRT-PCR assays confirmed that both *ECE1* and *EED1* are Efg1-regulated genes in the urine microenvironment (**Fig. 2**). Ece1, or candidalysin, is a known toxin required for mucosal and invasive infection, neutrophil recruitment, modulation of neutrophil extracellular traps, and immune defense (30, 39–41). Eed1 is known to play a crucial role in hyphal cell elongation and maintenance along with tissue dissemination (29, 38). To further understand the role of these factors during fungal CAUTI, we used deletion mutants (*ece1*Δ*/*Δ and *eed1*Δ*/*Δ) and overexpression mutants (*ECE1* OE and *EED1* OE) to assess their ability to form biofilms, morphology, and expression in our *in vitro* assays that mimic the catheterized bladder environment.

First, we assessed biofilm formation in Fg-urine conditions. We found that *ece1*Δ*/*Δ showed significant reduction of biofilm formation when compared to the WT (**Fig. 4A**). Overexpression of *ECE1* failed to restore the biofilm defect (**Fig. 4A**). Notably, the *eed1*Δ*/*Δ strain exhibited a significant biofilm deficiency, resembling the *efg1*Δ*/*Δ strains in both the SN250 and SC5314 backgrounds (**Fig. 4A**). However, overexpression of *EED1* partially restored biofilm formation (**Fig. 4A**).

**Figure 4.**
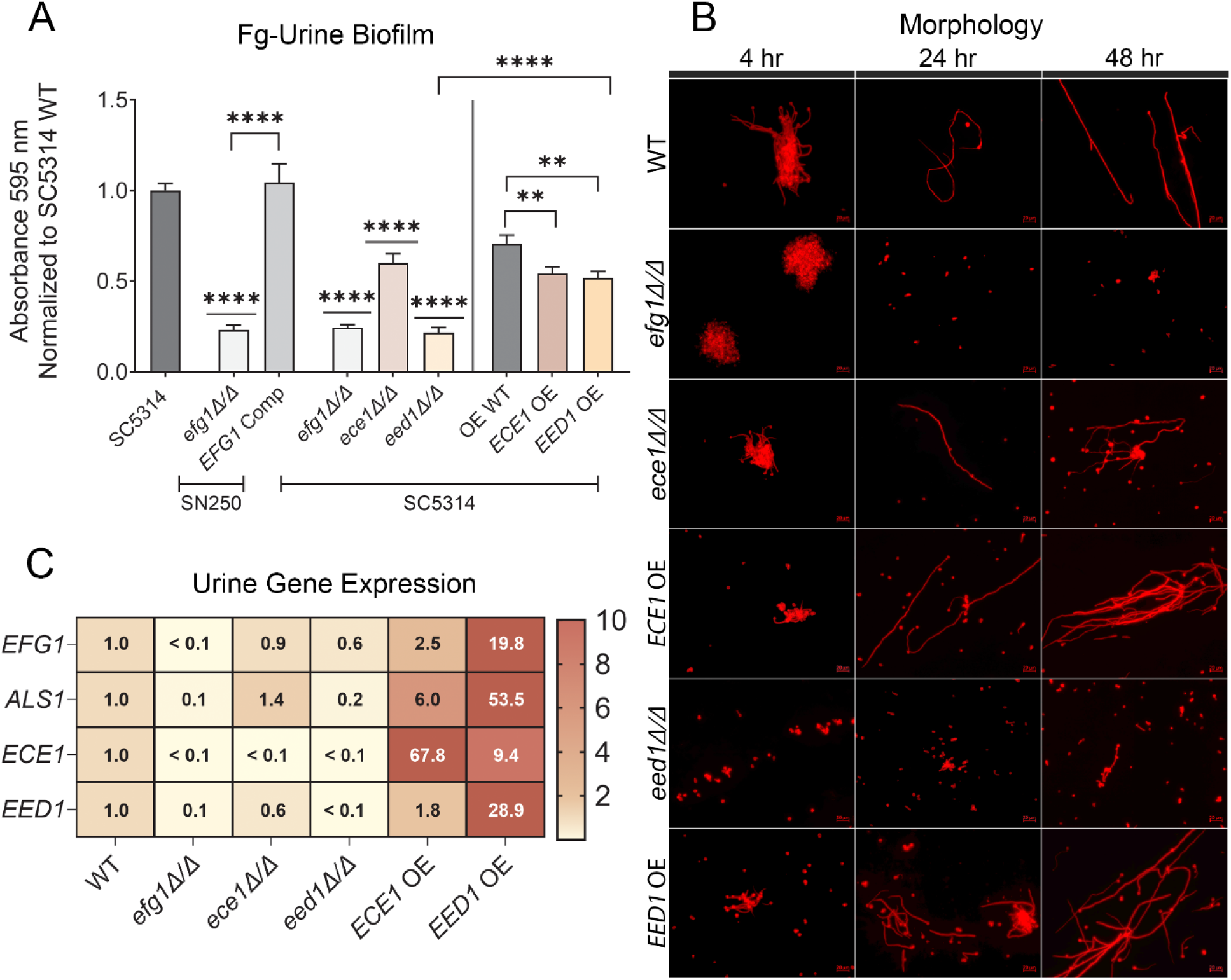
*In vitro* characterization of Ece1 and Eed1 mutants in urine conditions. (**A**) Forty-eight–hour Fg-dependent urine biofilm formation analysis of: *C. albicans* SN250 *efg1* Δ*/*Δ and *EFG1* complement strain; *C. albicans* SC5314 WT and deletion and overexpression mutants in the SC5314 background: *efg1* Δ*/*Δ, *ece1* Δ*/*Δ, *eed1* Δ*/*Δ, *ECE1* OE, and *EED1* OE. OE WT is the wildtype control strain for the overexpression mutants. Biofilm biomass was measured with crystal violet and the absorbance values at 595 nm were normalized to SC5314 WT. (**B**) SC5314 WT, SN250 *efg1*Δ*/*Δ, SC5314 *ece1*Δ*/*Δ, SC5314 *ECE1* OE, SC5314 *eed1* Δ*/*Δ, and SC5314 *EED1* OE morphologies were evaluated after 4, 24, and 48 hours of growth in urine + 10% serum. Representative images were taken at ×40 magnification following staining with calcofluor white (*Candida* cells; red). Scale bars, 20 µm. (**C**) *EFG1*, *ECE1*, *EED1*, and *ALS1* gene expression of RNA extracted from SC5314 WT, SN250 *efg1*Δ*/*Δ, SC5314 *ece1*Δ*/*Δ, SC5314 *ECE1* OE, SC5314 *eed1* Δ*/*Δ, and SC5314 *EED1* OE cells grown in urine for 48 hours. Gene expression is normalized to *ITS2* WT expression. Differences between groups for biofilm formation were tested for significance using the Mann-Whitney *U* test. Values represent means ± SEM derived from at least three independent experiments \**P* < 0.05, *\*\*P* < 0.005, \*\*\**P* < 0.0005, and \*\*\*\**P* < 0.0001.

Our morphology assessment in urine conditions showed that WT cells gradually filamented over 48 hours, while the *efg1*Δ*/*Δ cells remained yeast locked. The *ece1*Δ*/*Δ strain and its ECE1 OE counterpart were capable of filamenting in urine at 4, 24, and 48 hours, although the *ECE1* OE strain consisted primarily of filamentous cells at 48 hours (**Fig. 4B**). The *eed1*Δ*/*Δ strain showed predominately yeast and pseudohyphae cells at all times. While *eed1*Δ*/*Δ cells did not form full-length hyphae, long and short pseudohyphae cells were observed at 24- and 48-hours (**Fig. 4B**). In contrast, the *EED1* OE strain displayed hyperfilamentous cell morphologies, with the majority of cells being hyphal at 48 hours (**Fig. 4B**).

Since *EFG1* and *ALS1* are critical for biofilm formation and CAUTI, we evaluated their expression in the *ECE1* and *EED1* deletion mutants and overexpression counterparts. All strains were grown in urine conditions for 48 hours using WT strain as a reference and *efg1*Δ*/*Δ as negative control. RNA extraction was performed and the expression of *EFG1*, *ALS1, ECE1,* and *EED1* was quantified (**Fig. 4C**). In the *ece1*Δ*/*Δ strain*, EFG1, ALS1,* and *EED1* expression levels were comparable to WT while *ECE1* OE strain showed a greater than 2-fold increase in the expression of *ALS1* and *EFG1* (**Fig. 4C**). The *eed1*Δ*/*Δ strain exhibited *EFG1* expression levels similar to WT; however, it displayed a significant decrease of *ALS1* and *ECE1* expression. In contrast, the *EED1* OE showed a significant upregulation of all four factors (**Fig. 4C**). These data indicate that both *ALS1* and *ECE1* are downstream of *EED1* and that *EED1* levels also influence *EFG1* expression. Importantly, the *ALS1* downregulation in the *eed1*Δ*/*Δ strain provides an explanation for the significant biofilm deficiency in the Fg-urine biofilm assay (**Fig. 4A**), since Als1 is the main Fg-binding adhesin in this model.

### Ece1 and Eed1 are expendable factors for CAUTI establishment

We further evaluated the role of Ece1 and Eed1 in our mouse model of CAUTI by using the mutants and overexpression strains. Mice were transurethrally implanted with a silicone catheter and infected with ∼10^6^ colony forming units of the respective *C. albicans* strain. After 24 hours, mice were euthanized, and their organs and catheter were harvested for fungal burden assessment. We found that both *ece1*Δ*/*Δ and *eed1*Δ*/*Δ strains exhibited bladder and catheter colonization similar to WT (**Fig. 5A and B**). Notably, both *ECE1 and EED1* overexpression strains displayed fungal bladder and catheter burdens that did not differ from the WT (**Fig. 5A and B**), suggesting that overexpression of these proteins does not enhance infection. The OE WT control strain was also tested and showed a similar colonization profile to the SC5314 parent strain (**Fig. S5**). However, it was noted that a subgroup of OE infected mice had deficient bladder and catheter colonization compared to other OE infected mice (**Fig. 5A and B**). Due to the hyperfilamentous nature of these strains (**Fig. 4B**), bladder CFUs were also quantified via qPCR (**Fig. 5E, Fig. S6**) as the sticky, multinucleated hyphal cells may present as a singular colony when enumerating by serial dilution. Using qPCR to quantify fungal burden, bladder colonization of the OE mutants was found to be comparable to the deletion mutants, indicating that final fungal burden was underestimated when quantified by serial dilutions (**Fig. 5E**, **Fig. S6**). The *ECE1* OE and *eed1*Δ*/*Δ strains showed dissemination to the kidneys, but not as consistently as WT infected mice (**Fig. 5C**). Interestingly, despite their increased *in vitro* filamentation and biofilm formation (**Fig. 4A**), neither the *EED1* OE nor the *ECE1* OE strains showed any difference from WT in bladder colonization, catheter colonization, or systemic dissemination (**Fig. 5A-C**). This suggests that that neither an increase nor a complete loss of these factors provides a fitness advantage during fungal CAUTI (**Fig. 5A-C**).

**Figure 5.**
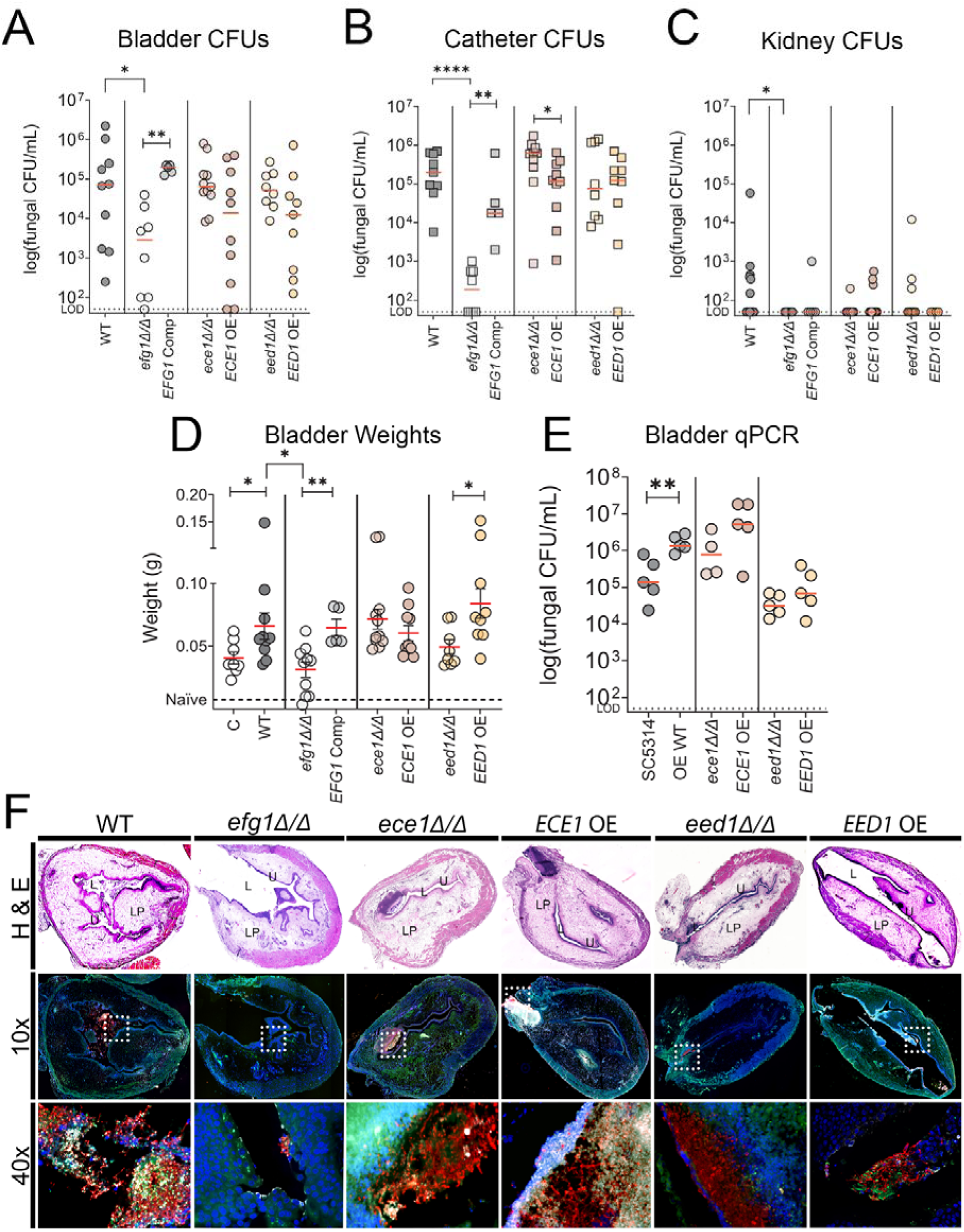
Ece1 and Eed1 are dispensable for fungal infection in the catheterized bladder environment. SC5314 WT, SN250 *efg1*Δ*/*Δ, SN250 *EFG1* complement, SC5314 *ece1*Δ*/*Δ, SC5314 *ECE1* OE, SC5314 *eed1*Δ*/*Δ, and SC5314 *EED1* OE fungal burden in (**A**) bladder, (**B**) catheter, (**C**) kidneys at 24 hours post infection and catheterization were quantitated as the number of CFUs recovered. For CFU enumeration, infections were done in at least two independent experiments with *n* = 3 or 4 mice for each one, and data are shown as the log(fungal CFU/organ or catheter). Animals that lost the catheter were not included in this work. (**D**) Bladder weights of naïve (non-implanted control; dashed line) mice, catheterized-only mice (‘C’), and mice catheterized and infected with *C. albicans* SC5314, SN250 *efg1*Δ*/*Δ, SN250 *EFG1* complement, SC5314 *ece1*Δ*/*Δ, SC5314 *ECE1* OE, SC5314 *eed1*Δ*/*Δ, or SC5314 *EED1* OE following 24h. Infections were done in two independent experiments with *n* = 3 or 4 mice for each one. (**E**) Catheterized bladder CFUs enumerated by qPCR for mice catheterized and infected with SC5314, OE WT, SC5314 *ece1*Δ*/*Δ, SC5314 *ECE1* OE, SC5314 *eed1*Δ*/*Δ, or SC5314 *EED1* OE. (**F**) Implanted and infected bladders and catheters were recovered at 24 hpi. Bladders were subjected to analysis by H&E (10x magnification) and IF staining (10x and 40x magnification). For the IF analysis, antibody staining was used to detect Fg (anti-Fg; green), *C. albicans* (anti-*Candida*; red), and neutrophils (anti-Ly6G; white). Staining with 4′,6-diamidino-2-phenylindole (DAPI; blue) delineated the urothelium and cell nuclei (representative images). Bladder lumen (L), urothelium (U), lamina propria (LP). White squares represent zoomed-in areas of higher magnification. The Mann-Whitney *U* test was used; *P* < 0.05 was considered statistically significant. \**P* < 0.05, \*\**P* < 0.005, \*\*\**P* < 0.0005, and \*\*\*\**P* < 0.0001. The horizontal dotted line represents the limit of detection (LOD) of viable fungi for (**A**), (**B**), (**C**), and (**E**). The horizontal red bar represents the median value.

Since catheterization induces bladder inflammation, which is then exacerbated by infection (42, 43), we measured bladder weights as a proxy for this inflammation response. WT-infected and catheterized bladders exhibited a significant increase of bladder inflammation when compared to catheterized-only and naïve mouse bladders (**Fig. 5D**). The *efg1*Δ*/*Δ *-*catheterized and infected bladders were similar in weight to catheterized-only bladders, indicating the mutant did not contribute to inflammation (**Fig. 5D**). The *ece1*Δ*/*Δ and *eed1*Δ*/*Δ -catheterized and infected bladders did not differ statistically from WT (**Fig. 5D**). Although no change in bladder weight was observed between *ECE1* OE bladders and *ece1*Δ*/*Δ or WT, *EED1* OE infected had significantly higher weights than their *eed1*Δ*/*Δ counterparts. The fact that *EED1* OE induces hyperfilamentous cells (**Fig. 4B**) may be responsible for this since *Candida* filamentation induces a stronger inflammatory immune response (44, 45).

This prompted us to investigate the host-fungal interactions within the catheterized bladder using H&E and immunofluorescence (IF) staining (**Fig. 5F**). In WT infected bladders, yeast and hyphal cells formed with Fg-associated biofilms in the lumen and penetrated the urothelium despite neutrophil recruitment (**Fig. 5F** and **Fig. S7,8** for single channels). In sharp contrast, the *efg1*Δ*/*Δ-infected bladder showed negligible fungal burden, with only small yeast cell aggregates (**Fig. 5F** and **Fig.S7,8** for single channels). Bladders infected with *ece1*Δ*/*Δ or *ECE1 OE* contained yeast and hyphal cells interacting with Fg and immune cells (**Fig. 5F** and **Fig.S7,8** for single channels). Notably, both showed invasive hyphae entering the urothelium and encountering a strong immune response (**Fig. 5F** and **Fig.S7,8** for single channels).

The *eed1*Δ*/*Δ infected bladder showed aggregated/clustered yeast and pseudohyphae cells in the bladder lumen, penetrating the urothelium and encountering immune cells (**Fig. 5F** and **Fig. S7,8** for single channels). Interestingly, the *EED1* OE bladder showed isolated biofilms comprised mostly of hyphal cells colocalizing with Fg and interacting with immune cells (**Fig. 5F** and **Fig. S7,8** for single channels). Overall, the imaging revealed stark differences in fungal burden and immune cell infiltration in the catheterized bladders among the fungal strains. Despite these histological differences between the *ECE1* and *EED1* strains, the CFU enumeration suggest that Ece1 and Eed1 are dispensable factors for colonization of the catheterized bladder environment (**Fig. 5**).

## DISCUSSION

*C. albicans* is a leading CAUTI pathogen, yet its molecular pathogenesis in this environment is poorly understood. We recently identified Efg1 as a key driver of *C. albicans* CAUTIs (11). Although Efg1 is a known master virulence regulator, its regulon is both environment-dependent and variable across clinical strains, even under identical conditions (22, 23). Given the bladder’s unique, dynamic, and hostile environment (hypertonic, acidic, and urea-rich) (16, 46, 47), we sought to define the urine-specific Efg1 regulon of *C. albicans*.

Here, we identified the SC5314 Efg1 urine-specific regulon and used it to start dissecting fungal CAUTI pathogenesis. This regulon includes novel Efg1-regulated associations, such as 26 previously uncharacterized genes (**Fig. 2C, Table S9**). Recent studies have found that gene functions identified in the standard lab strain SC5314 often do not apply to clinical isolates; rather, many functions are strain-dependent (31, 48). For example, Wakade *et al* 2024 found that SC5314 *nrg1*Δ*/*Δ forms filamentous cells, but *NRG1* deletions in some clinical strain backgrounds were only capable of forming pseudohyphae, indicating that Nrg1 function is dependent on the genetic background of the strain (31). Furthermore, one clinical isolate with an *NRG1* deletion was found to have increased expression levels of *ECE1 in vitro* but decreased *ECE1* expression in *in vivo* assays (31). Another study found that a loss of function *EFG1* mutation in a clinical isolate from a bloodstream infection (P94015) led to increased fitness in infection models despite reduced filamentation, contrary to the decreased virulence seen in *EFG1* deletions in many other clinical and laboratory strains (48). These studies show the complexity of clinical isolates and the need to use and study clinical strains in the laboratory to better understand the plasticity of human fungal disease.

We previously found that Efg1 was critical for several clinical *C. albicans* urinary isolates to cause CAUTI in our mouse model, again demonstrating its importance for causing this infection. However, crucial evidence came from our patient data; fungal transcriptomics from 9 patient catheters showed upregulation of the same Efg1-urine factors identified in our Fg-urine assay (**Fig. 2 and 3**). Importantly, the Efg1 transcriptional profiles across all 9 catheters showed ∼80% similarity (**Fig. 3C**), highlighting this network’s importance during human CAUTI. This data concludes that Efg1-urine network is clinically relevant and corresponds to human infections. Accordingly, clinical isolates must continue to be studied alongside lab-domesticated strains to ensure a comprehensive understanding of *C. albicans* pathogenesis, accounting for both strain variation and environmental context.

Based on their expression profiles *in vitro* and during human infection, we selected *ECE1* and *EED1* to characterize their roles in CAUTI using deletion and overexpression strains. We evaluated their ability to form Fg-urine biofilms, filament in urine, and cause infection in our CAUTI mouse model. Despite these distinct *in vitro* phenotypes, both deletion and overexpression mutants of *ECE1* and *EED1* colonized the bladder and catheter as effectively as WT in our mouse model (**Fig. 5**). This suggests that *ECE1* and *EED1* are ultimately dispensable factors during fungal CAUTI, at least in a short-term model. This finding was puzzling, given both the observed fungal bladder tissue invasion (**Fig. 5F**) and the filament-inducing nature of the catheterized bladder environment (**Fig. 4B**)(11).

The dispensability of *ECE1* was particularly surprising, as its product, candidalysin, is critical for host epithelial damage and tissue invasion (49–51). However, recent studies shed light on why this might be the case, as candidalysin is neutralized by albumin (52, 53). This is highly relevant, as catheter-induced inflammation promotes protein extravasation, making serum albumin and Fg two of the most abundant proteins in the catheterized bladder (10, 12, 35, 42). Consequently, the candidalysin toxin is neutralized in this albumin-rich environment, suggesting that the catheterized environment inadvertently mitigates exacerbated tissue damage and the full pathogenic potential of *C. albicans* during CAUTI. Interestingly, hypoalbuminemia (low serum albumin) is linked to increased risk and severity of fungal infections, including candidemia and pneumonia, and correlates with poorer clinical outcomes (54, 55). This association is also observed in kidney transplant patients (56), who are catheterized pre- and post-surgery (57), where co-occurring UTIs lead to particularly poor outcomes (56). Therefore, catheterized patients with hypoalbuminemia may be highly susceptible to severe CAUTI and its complications, especially fungal CAUTI.

Since candidalysin is a secreted factor (51), its activity may also be mitigated by the bladder’s urodynamics. The constant flushing of urine and urine production inherently reduces the toxin’s overall concentration, preventing it from reaching potent or effective levels (58–60). The fact that candidalysin does not contribute to fungal CAUTI highlights the need to study fungal infections in their context-specific environment to truly understand virulence and pathogenesis mechanisms of varying infection types. Furthermore, this work highlights the need to understand the contribution of other Efg1-regulated factors.

The ability of the *eed1*Δ*/*Δ mutant to colonize the catheterized bladder was a surprising result, given its deficient *in vitro* Fg-urine biofilm formation (**Fig. 4A**). This *in vivo* success may be explained by the interplay between the mutant’s traits and the host environment. The *EED1* mutant is characterized by rapid proliferation (forming yeast and pseudohyphae) (38), while the urinary catheter itself causes urothelial damage and plasma extravasation, creating a continuously nutrient-rich environment (7, 9, 10, 33, 35, 61). Therefore, the mutant’s rapid proliferation is likely a significant advantage in this nutrient-replete setting, compensating for its biofilm deficiency. This aligns with previous studies, which have shown that despite the *eed1*Δ/Δ mutant’s inability to maintain true hyphal cell types, it is still virulent in a mouse systemic model of infection (29, 38). We propose that during CAUTI, this mutant’s morphological flexibility (shifting from yeast to pseudohyphae) followed by rapid proliferation is sufficient to damage the bladder urothelium and allow for colonization. In support of this, our bladder imaging shows *eed1*Δ/Δ fungal cell aggregates breaching the urothelium (**Fig. 5F**). While the majority of these cells appear to be in yeast form, the presence of pseudohyphae appears sufficient for tissue invasion. For these aggregates to form and colonize, however, adequate adhesion is also required. *ALS1* expression is lacking in this strain (**Fig. 4C**), suggesting the fungal cells may be employing a different adhesin for fungus-fungus or fungus-urothelium adhesion. Future studies identifying other urine-specific adhesins could be beneficial in uncovering additional fungal-urine interactions.

Another critical observation arising from our study is the shift and variation in *C. albicans’* transcriptional profile of human urinary catheters in the presence of other microbes. This aligns with previous studies showing that multi-species interactions cause global transcriptional changes in *C. albicans,* specifically with *Streptococcus gordonii, Fusobacterium nucleatum, Enterococcus faecalis, S. aureus,* and *Streptococcus mutans* (*62–65*). In our cohort, this was exemplified by catheter HUC103, which was colonized solely by *C. albicans* and showed the highest expression of *EFG1*, *ALS1*, *EED1*, and *ECE1* (**Fig. 3**). In contrast, all other catheters were polymicrobial (containing 1-4 other pathogens) and, while these genes were still upregulated, their expression levels were lower than in the mono-microbial HUC103 sample. Despite this observed variability in the Efg1 regulon, the Efg1 virulence networks were highly conserved, showing ∼80% similarity across all 9 catheters. This “variability within conservation” suggests two possibilities, the differences stem from strain-specific factors, or (equally likely) the Efg1-urine regulon is directly modulated by co-existing microbes. Given that ∼77% of CAUTIs are polymicrobial, this result warrants further dissection of fungal-uropathogen responses (66–68).

The finding that Efg1’s importance is conserved across diverse clinical isolates, including polymicrobial infections, is critical. This high degree of conservation confirms its clinical applicability, making Efg1 and its downstream effectors promising therapeutic targets for *C. albicans* CAUTIs.

This study establishes the Efg1-urine regulon as a clinically conserved and central driver of human *C. albicans* CAUTI. By demonstrating that this virulence network is highly similar (∼80%) across diverse, polymicrobial patient catheter infections, this work overcomes the challenge of strain-to-strain variability. It validates the Efg1 network as a high-priority, clinically applicable therapeutic target, even while revealing that canonical virulence factors like candidalysin are surprisingly dispensable in the unique, albumin-rich environment of the catheterized bladder.

## Materials and Methods

### Ethics statement

All animal care was consistent with the Guide for the Care and Use of Laboratory Animals from the National Research Council. The University of Notre Dame Institutional Animal Care and Use Committee approved all mouse infections and procedures as part of protocol number 25-01-9009. For urine and blood collections, all donors signed an informed consent form, and protocols were approved by the Institutional Review Board of the University of Notre Dame under study #19-04-5273 for urine and #18-08-4834 for blood.

### Urine collection

Human urine from at least two healthy female donors between the ages of 20 to 35 were collected and pooled. Donors did not have a history of kidney disease, diabetes, or recent antibiotic treatment. Urine was sterilized with a 0.22-μm filter (VWR 29186-212), pH was normalized to 6.0 to 6.5, and urine was used immediately for the assays.

### Collection of human urinary catheters

Patient catheters were collected with informed consent after the clinical decision to remove for standard of care was made as described (66). This study was approved by the Washington University School of Medicine (WUSM) Internal Review Board (approval #201410058) and performed in accordance with WUSM’s ethical standards and the 1964 Helsinki declaration and its later amendments.

### Fungal cultures

All strains used in this study are in SC5314 or in SN250, a derivative of the SC5314 background strain used to generate Suzanne Noble’s library (26), and are listed in **Table S11**. All strains were cultured at 37°C with aeration in 5 ml of YPD [yeast extract (10 g/liter; VWR J850-500G), peptone (20 g/liter; VWR J636-500G), and dextrose (20 g/liter; VWR BDH9230-500G)] broth. For *in vivo* mouse experiments, *C. albicans* strains were grown static overnight (to simulate gut conditions, the presumed source in clinical infections) in 10 ml of YPD. Most assays were done in SC5314, but for the RNA-sequencing we needed to use the SN250 *efg1*Δ*/*Δ and its corresponding *EFG1* complement as the *EFG1* complement strain was necessary for appropriate analysis. Our current CRISPR methods do not make it feasible to produce a complement or overexpression *EFG1* strain in the SC5314 background; thus, we used the SN250 strains for RNA-sequencing and validated our findings in both SN250 *efg1*Δ*/*Δ and SC5314 *efg1*Δ*/*Δ.

### Generation of CRISPR-Cas9 Deletion Strains

The CRISPR-Cas9 plasmid for the *ECE1* gene deletion was constructed and transformed into *C. albicans* as previously described with some modifications (69, 70). Briefly, the *ECE1*-targeting gene drive was ordered as a gene fragment from Genscript and cloned into the backbone plasmid (Addgene #89576) via Gibson Assembly following digestion with NgoMIV. Plasmids were PCR verified for the presence of the insert and co-transformed into *C. albicans* cells along with ∼60ng of the gene fragment (ie. repair template) into NEUT5L (neutral locus) following the lithium acetate-based yeast transformation protocol (70). Transformants were PCR-verified for the deletion by comparing to the non-transformed WT strain.

### Generation of CRISPR Overexpression Strains

To generate the overexpression strains, we used the *C. albicans* hyperdCas12a CRISPR activation system (71). The crRNAs were designed with the Eukaryotic Pathogen CRISPR guide RNA/DNA Design Tool with a custom 5’ “TTTV” PAM and with the C. albicans SC5314 FungiDB-26 genome selected (72), and cloned into pRS1005 (AddGene ID: 247641) as previously described (71). Briefly, forward and reverse-complement DNA oligo pairs corresponding to each array were synthesized by Integrated DNA Technologies (IDT), resuspended in Nuclease Free Duplex Buffer from IDT (cat. 11-05-01-12) to 100μM, incubated at 94°C for 1 min, mixed with the counterpart oligo in an equal volume, and incubated at 94°C for 2 min. The duplexed oligo pairs were then cloned into the *C. albicans* hyperdCas12a CRISPRa plasmid via a Golden Gate strategy using ∼1000ng of plasmid, 1μL of duplexed oligo, 2μL of 10X rCutSmart buffer, 2μL of ATP, 1μL of SapI, 1μL of T4 DNA ligase, and nuclease-free water to a total volume of 20μL. Cloning mixes were then incubated in a thermocycler using the following conditions: (37°C for 2 min, and 16°C for 5 min) for 99 cycles, 65°C for 15 min, and 80°C for 15 min. The next morning, 1μL of SapI was added to each reaction and incubated at 37°C for 1h to remove any leftover uncloned plasmids. Plasmids were PCR tested for the presence of the correct crRNA before proceeding, as previously described (73). Plasmids were then linearized with the PacI enzyme from NEB (cat R0547L), left at 37°C for 12h, 65°C for 20 min, and transformed into C. albicans with a lithium acetate method as previously described (71). Briefly, pelleted overnight cultures of C. albicans were incubated for 1h with a mix containing 800μL of 50% polyethylene glycol (PEG), 100μL of 10X Tris-EDTA buffersolution, 100μL of 1M lithium acetate, 40μL of UltraPureTM Salmon Sperm DNA Solution from ThermoFisher (cat. 15632-011), and 20μL of 1M dithiothreitol (DTT). Cell solutions were then incubated at 42°C for 50 min in a water bath, pelleted, and washed three times with 1mL of YPD. The cells were then diluted to a final volume of 10mL in YPD and incubated at 30°C for 4h (250 RPM). Pelleted and resuspended cells were then plated onto YPD + NAT (250μg/mL) and incubated statically at 30°C for 2 days. Individual transformants were then patched onto YPD + NAT and grown statically at 30°C for an additional day. Transformants were PCR-tested for the correct crRNA as well as for successful integration into the NEUT5L region, as previously described [3]. crRNA Sequences for hyperdCas12a CRISPRa EED1 (DEF1) were F-cgataatttctactaagtgtagatGGTATATAAATAGACAGAAT and R-aacATTCTGTCTATTTATATACCatctacacttagtagaaatta. The crRNA sequences used for the hyperdCas12a CRISPRa ECE1 strain were F-cgataatttctactaagtgtagatCTGATCCTATTGGGTGCAAT and R-aacATTGCACCCAATAGGATCAGatctacacttagtagaaatta.

### Biofilm formation assays

Biofilm formation assays were performed in 96-well flat-bottomed plates (VWR, 10861-562) coated with 100 μl of Fg (150 μg/ml) incubated overnight at 4°C. The various strains were grown as described above, and the inoculum was normalized to ∼1 × 10^6^ CFUs/ml. Cultures were then diluted (1:100) into human urine supplemented with 10% heat inactivated human serum and incubated in the wells of the 96-well plate at 37°C for 48 hours while static.

### Crystal violet staining

Following biofilm formation on Fg-coated microplate, the supernatant was removed, and the plate was incubated in 200 μl of 0.5% crystal violet for 15 min. Crystal violet stain was removed, and the plate was washed with water to remove the remaining stain. Plates were dried and then incubated with 200 μl of 33% acetic acid for 15 min. In another plate, 100 μl of the acetic acid solution was transferred, and absorbance values were measured via a plate spectrophotometer at 595 nm (Molecular Devices SpectraMax ABS). Values were normalized to SC5314 WT.

### Construction of protein-protein interaction networks and GO slim analysis

Using the Search Tool for the Retrieval of Interacting Genes (STRING) database by Cytoscape Software, functional interactions between factors that promoted Fg-urine biofilm formation, factors that resulted in defective Fg-urine biofilm formation, and factors found to comprise the Efg1-urine regulon were screened and mapped. The *Candida* Genome Database (candidagenome.org) Slim Mapper tool was used to categorize genes according to GO-Slim terms. Efg1-urine regulons were compared among WT-urine, *efg1Δ/Δ*-urine, WT-YPD, and *efg1Δ/Δ*-YPD regulons using an E Venn network (74) online software (http://www.ehbio.com/test/venn/#/) to visualize the uniquely Efg1 regulon in Fg-urine biofilms.

### RNA extraction

Total RNA was extracted from biofilms with the Zymogen RNA extraction kit (Zymo Research catalog no. R2071), and subsequently DNase I–treated (Thermofisher scientific catalog no. 89836).

### RNA sequencing and bioinformatics

Extracted RNA was sent to Azenta Life Science for RNA sequencing and bioinformatics analysis. Whole transcriptome sequencing was done following rRNA depletion. Sequence reads were trimmed to remove possible adapter sequences and nucleotides with poor quality using Trimmomatic v.0.36. The trimmed reads were mapped to the *Candida_albicans*_30-877911309 reference genome available on ENSEMBL using the STAR aligner v.2.5.2b. The STAR aligner is a splice aligner that detects splice junctions and incorporates them to help align the entire read sequences. BAM files were generated as a result of this step. Unique gene hit counts were calculated by using featureCounts from the Subread package v.1.5.2. The hit counts were summarized and reported using the gene_id feature in the annotation file. Only unique reads that fell within exon regions were counted. After extraction of gene hit counts, the gene hit counts table was used for downstream differential expression analysis. Using DESeq2, a comparison of gene expression between the customer-defined groups of samples was performed. The Wald test was used to generate p-values and log2 fold changes. Genes with an absolute log2 fold change > 2 were identified as differentially expressed genes for each comparison.

For the patient catheter RNA, the total of the gene hit counts for each sample was summed. For each gene in every sample, counts per million (CPM) were calculated using the following formula: (Gene Hit Count/Total Gene Hit Counts of Sample) *10^6^. The counts per million of each gene were then normalized to the counts per million value of the housekeeping gene ARP3 for each sample. Expression values are graphed as CPM normalized to ARP3.

### Morphology assessment

All strains of *C. albicans* were grown in human urine with 10% serum. At 4, 24, and 48 hours, a sample of each condition was taken, fixed with 10% formalin, and stained with calcofluor (100 μg/ml). Samples were viewed under a Zeiss inverted light microscope (Carl Zeiss, Thornwood, NY) with the 4′,6-diamidino-2-phenylindole fluorescent channel. Representative images were taken at 40× magnification and processed with ImageJ.

### *In vivo* mouse model

Mice used in this study were ∼6-week-old female WT C57BL/6 mice purchased from the Jackson Laboratory. Mice were subjected to transurethral implantation of a silicone catheter and inoculated as previously described (75). Briefly, mice were anesthetized by inhalation of isoflurane and implanted with a 6-mm-long silicone catheter (Braintree Scientific, SIL 025). Mice were infected immediately following catheter implantation with 50 μl of ∼1 × 10^6^ CFUs/ml in PBS of one of the fungal strains introduced into the bladder lumen by transurethral inoculation. Mice were euthanized at 24 hpi by cervical dislocation after anesthesia inhalation, and the catheter, bladder, kidneys, spleen, and heart were aseptically harvested. Bladders were weighed and then homogenized for CFUs or fixed in 10% formalin overnight before being paraffin-embedded, sectioned, stained, and imaged. The other organs were homogenized, and catheters were cut into small pieces before sonication for fungal CFU enumeration.

### Immunohistochemistry and H&E staining of mouse bladders

Mouse bladders were fixed in 10% formalin overnight, before being processed for sectioning and staining as previously described (33). Briefly, bladder sections were deparaffinized, rehydrated, and rinsed with water. Antigen retrieval was accomplished by boiling the samples in Na-citrate, washing in tap water, and then incubating in 1× PBS three times. Sections were then blocked [1% BSA and 0.3% Triton X-100 (Acros Organics, 21568-2500) in 1× PBS], washed in 1× PBS, and incubated with appropriate primary antibodies diluted in blocking buffer overnight at 4°C. Next, sections were washed with 1× PBS, incubated with secondary antibodies for 2 hours at room temperature, and washed once more in 1× PBS before Hoechst dye staining. Secondary antibodies for immunohistochemistry were Alexa Fluor 488 donkey anti-goat, Alexa Fluor 550 donkey anti-rabbit, and Alexa Fluor 650 donkey anti-rat. H&E stain for light microscopy was done by the core facilities at the University of Notre Dame (ND Integrated Imaging core). All imaging was done using a Zeiss inverted light microscope. Zen Pro, ImageJ software, and IMARIS Image Analysis software were used to analyze the images.

### qRT-PCR

RNA was reverse transcribed into cDNA via qScript cDNA synthesis kit (QuantaBio, catalog no. 101414); incubated for 5 min at 25°C, 30 min at 42°C, and 5 min at 85°C; and held at 4°C. With FAST Start Sybr Green mastermix, qPCR was performed on CFX Opus 96 (BioRad) under the following conditions: 95°C^5min^ (95^15sec^, 60^60sec^)_35 cycles_ (95^15sec^, 60^60sec^, 95^15sec^) _melt curve_. Data were normalized using *ITS2* (internal transcribed spacer 2) as an internal housekeeping control gene and WT as a sample calibrator with the expression set to 100% or 1. Data were analyzed via the ΔΔCt method. Primers are described in **Table S12**.

For quantification of bladder colonization, fungal gDNA was extracted from bladder samples and *C. albicans efg1*Δ*/*Δ culture using the Wizard genomic DNA extraction (Promega, catalog no. A1120). An 8 point, 1:10 serial dilution standard curve was made using the *C. albicans efg1*Δ*/*Δ culture. After gDNA isolation, qPCR was performed for bladder samples and *C. albicans efg1*Δ*/*Δ culture on CFX Opus 96 (BioRad) with Fast Start Sybr Green mastermix using the *ITS2* primers described in **Table S12**. The following thermocycler conditions were used: 95°C^10min^ (95^15sec^, 60^30sec^)_35 cycles_ (95^15sec^, 60^60sec^, 95^15sec^) _melt curve._ The CFX Maestro program was used for analysis.

### Statistical analysis and reproducibility

Data from at least three experiments were pooled for each assay. Two-tailed Mann-Whitney *U* tests were performed with GraphPad Prism 5 software (GraphPad Software, San Diego, CA) for all comparisons described in biofilm, and CAUTI mouse experiments. Values represent means ± SEM derived from at least three independent experiments (\**P* < 0.05; \*\**P* < 0.005; \*\*\**P* < 0.0005; \*\*\*\**P* < 0.0001; and ns, difference not significant).

Bray-Curtis multivariate analysis of similarity (ANOSIM) analysis was used to examine statistical significance among the fungal transcriptional profiles from catheterized patients with a fungal infection. The dendrogram and cluster analysis based on the transcriptional profile similarly matrix generated based on ANOSIM were generated using Primer-E 7.0 software.

Non-metric MDS plot was used to examine statistically significant differences among samples as a horizontal distance. An important component of the plot is a measure of the goodness of fit of the final plot, termed the “stress.” A stress greater than 0.2 indicates that the plot is close to random, a stress less than 0.2 indicates a useful two-dimensional picture, and a stress less than 0.1 corresponds to an ideal ordination with no real prospect of misinterpretation. Stress was calculated as described by Kruskal (32) with the Primer-E 7.0 software.

## Acknowledgements

We thank members of the laboratories of FHST and ALFM for helpful suggestions and insightful comments. We give special thanks to Sara Cole, Sarah Chapman, and the ND Integrated Imaging Facility for tissue processing and support during imaging.

## Author Contributions

FHST and ALFM conceived and supervised the research. AAL, NCG, KNK, ALFM, and FHST conducted the experiments and sample and data analyses. ALFM and CLPO conducted human sample collection. IDJ shared the *eed1Δ/Δ* strain. FHST, ALFM, and AAL wrote the manuscript. FHST, ALFM, AAL, KNK, NCG, RSS, IDJ, MGC, and SJH reviewed and edited the final manuscript.

