## Supplementary Material for "The Unique Efg1 Fungal Virulence Regulon in the Catheterized Bladder Environment"

**Supplementary Figures:**

**Fig. S1.** Transcriptional profile of *EFG1* complement relative to WT in Fg-urine and Fg-YPD biofilm conditions.

**Fig. S2.** Transcriptional profile of *efg1∆/∆* Fg-YPD relative to WT Fg-YPD biofilms.

**Fig. S3.** Validation of Efg1-urine regulon in SC5314 *efg1∆/∆* Fg-urine biofilms.

**Fig. S4.** *C. albicans* SC5314 and OE WT control bladder weights and fungal burden during CAUTI.

**Fig. S5.** Spleen and heart colonization during mouse CAUTI.

**Fig. S6.** Bladder colonization during CAUTI quantified by qPCR compared to CFUs.

**Fig. S7.** Single channels of *C. albicans* WT and mutants bladder colonization during CAUTI at 10x magnification.

**Fig S8.** Single channels of *C. albicans* WT and mutants bladder colonization during CAUTI at 40x magnification.

**Supplementary Tables:**

**Table S1.** STRING network protein-protein node interactions of factors that promoted biofilm formation.

**Table S2.** STRING network protein-protein node interactions of factors with defective biofilm formation.

**Table S3.** Gene ontology slim analysis of significant biofilm formers (decreased biofilm formation, promoted biofilm formation).

**Table S4.** WT v *efg1∆/∆* in Fg-urine biofilms RNA sequencing hits with log2FoldChange greater than the absolute value of 2.

**Table S5.** WT v *EFG1* complement in Fg-urine biofilms RNA sequencing hits with log2FoldChange greater than the absolute value of 2.

**Table S6.** WT v *EFG1* complement in Fg-YPD biofilms RNA sequencing hits with log2FoldChange greater than the absolute value of 2.

**Table S7.** WT Fg-YPD v *efg1∆/∆* Fg-YPD biofilms RNA sequencing hits with log2FoldChange greater than the absolute value of 2.

**Table S8.** Downregulated RNA sequencing hits in *efg1∆/∆* in urine and YPD when compared to WT.

**Table S9.** GO-slim ontology analysis of the Efg1-regulon in urine.

**Table 10.** STRING network protein-protein node interactions of Efg1-regulon in urine.

**Table S11.** Strains used in this study.

**Table S12.** Primers used in this study.

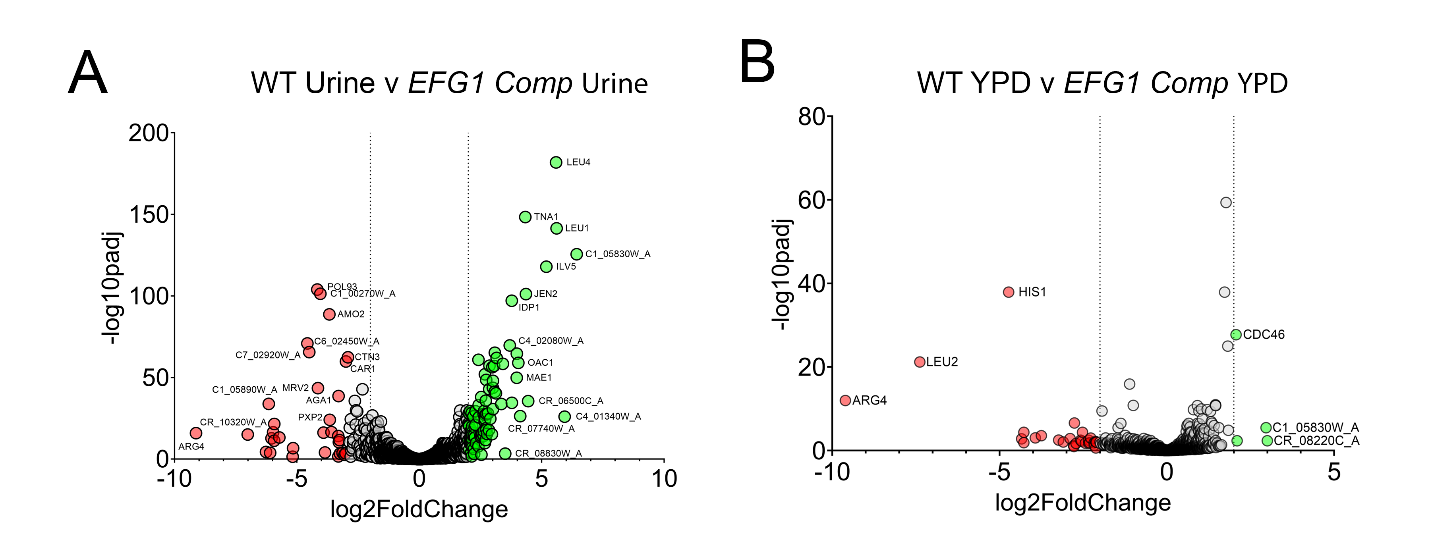

**Fig. S1.** **Transcriptional profile of *EFG1* complement relative to WT in Fg-urine and YPD biofilm conditions.** Volcano plot showing RNA sequencing results comparing *EFG1* complement (**A**) Fg-urine or (**B**) Fg-YPD biofilm cells to WT Fg-urine or YPD biofilm cells. Following whole genome sequencing, gene expression comparison between the *EFG1* complement and WT biofilms in urine or YPD was conducted using DESeq2. The Wald test was used to generate p-values and log2 fold changes. Genes with an absolute log2 fold change > 2 were identified as differentially expressed genes. Red indicates downregulation of *EFG1* complement compared to WT (log2FoldChange ≤ -2). Green indicates upregulation of *EFG1* complement compared to WT (log2FoldChange ≥ 2).

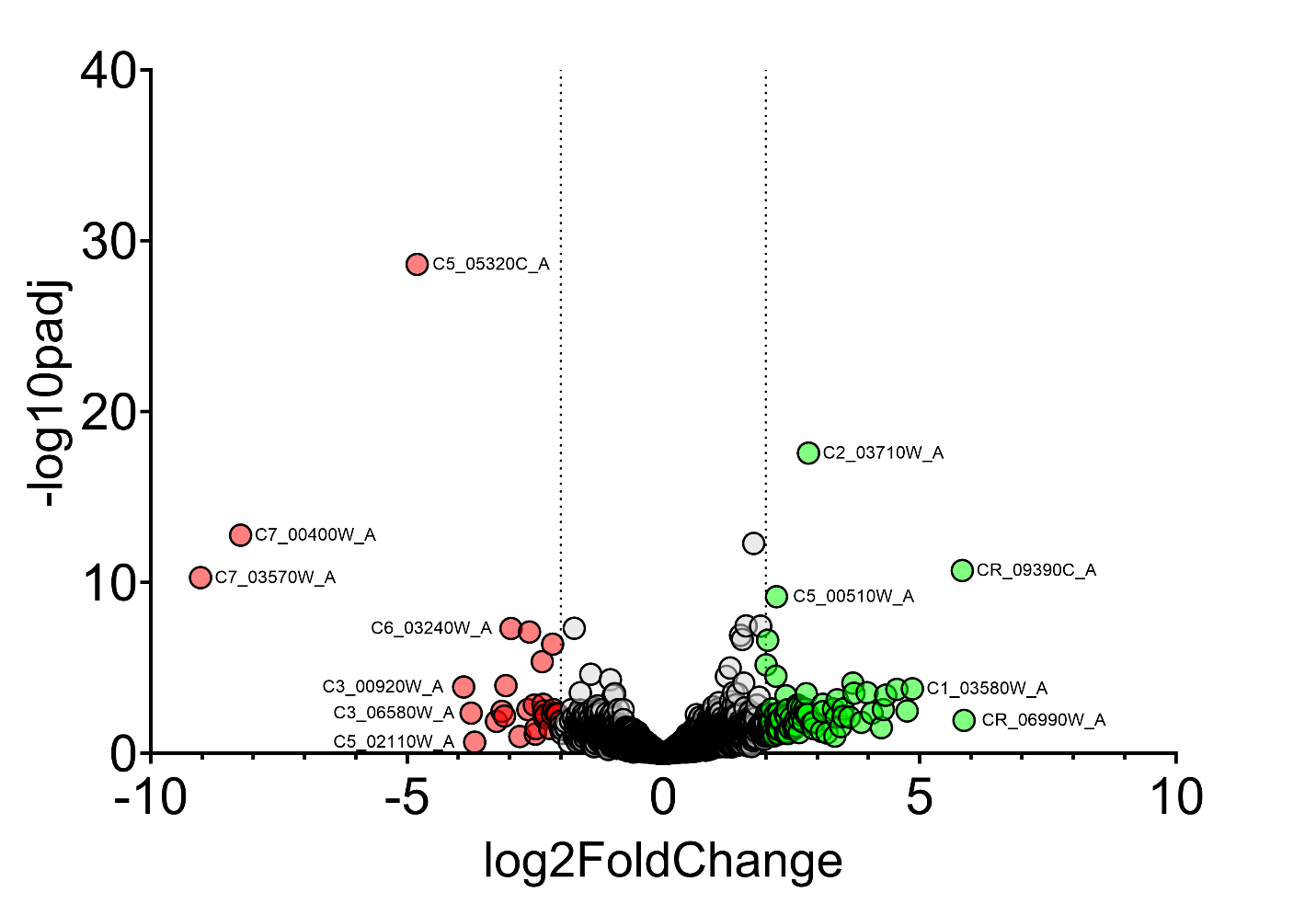

**Fig. S2. Transcriptional profile of *efg1∆/∆* Fg-YPD relative to WT Fg-YPD biofilms**. Volcano plot showing RNA sequencing results comparing *efg1∆/∆* Fg-YPD biofilm cells to WT Fg-YPD biofilm cells. Following whole genome sequencing, gene expression comparison between Fg-YPD biofilms was conducted using DESeq2. The Wald test was used to generate p-values and log2 fold changes. Genes with an absolute log2 fold change > 2 were identified as differentially expressed genes. Red indicates downregulation of *efg1∆/∆* YPD compared to WT YPD (log2FoldChange ≤ -2). Green indicates upregulation of *efg1∆/∆* YPD compared to WT YPD. (log2FoldChange ≥ 2).

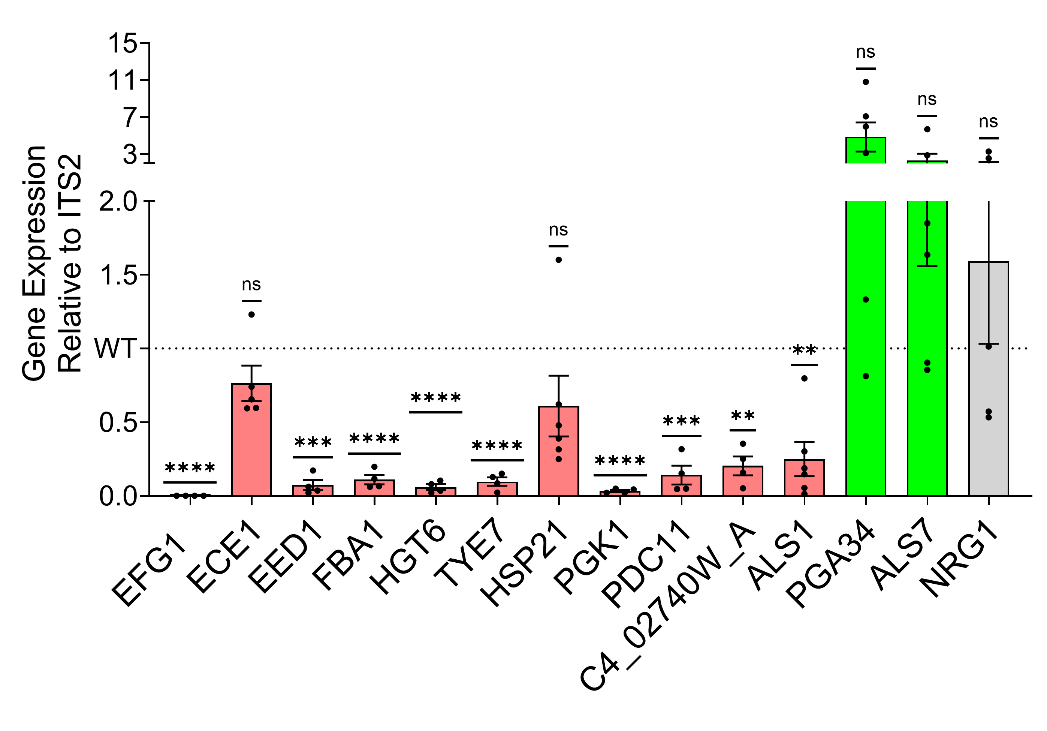

**Fig. S3.** **Validation of Efg1-urine regulon in SC5314 *efg1∆/∆* Fg-urine biofilms.** Validation of Efg1-urine regulon in SC5314 background via qRT-PCR. RNA was extracted from SC5314 WT and *efg1∆/∆* 48-hour Fg-urine biofilms, and expression of select Efg1-urine regulon genes were found by qRT-PCR and analyzed with the 2^-∆∆Ct^ method. Data are shown as the relative expression normalized to the housekeeping gene, *ITS2*, and SC5314 WT.

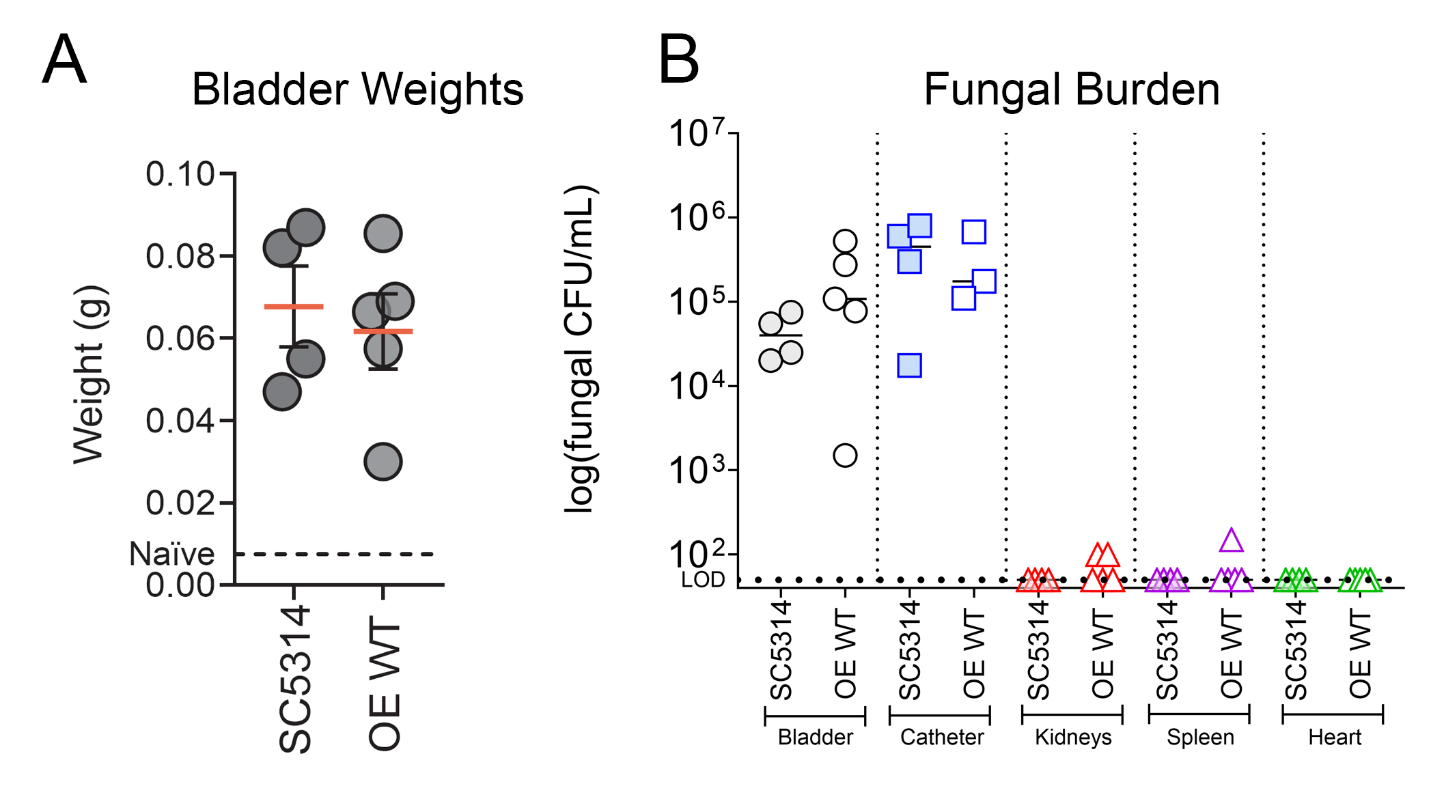
**Fig. S4. *C. albicans* SC5314 and OE WT control bladder weights and fungal burden during CAUTI. (A)** Bladder weights of naïve (non-implanted control; dashed line) mice and mice catheterized and infected with *C. albicans* SC5314 or OE WT after 24h. (**B**) Fungal burden of SC5314 and OE WT on the harvested organs and catheter after 24h. Infections were done in two independent experiments with *n* = 3 or 4 mice for each one. Data are shown as the log(fungal CFU/organ or catheter). Animals that lost the catheter were not included in this work. The Mann-Whitney *U* test was used; *P* < 0.05 was considered statistically significant. **P* < 0.05, ***P* < 0.005, ****P* < 0.0005, and *****P* < 0.0001. The horizontal dotted line represents the limit of detection (LOD) of viable fungi. The horizontal bar represents the median value.

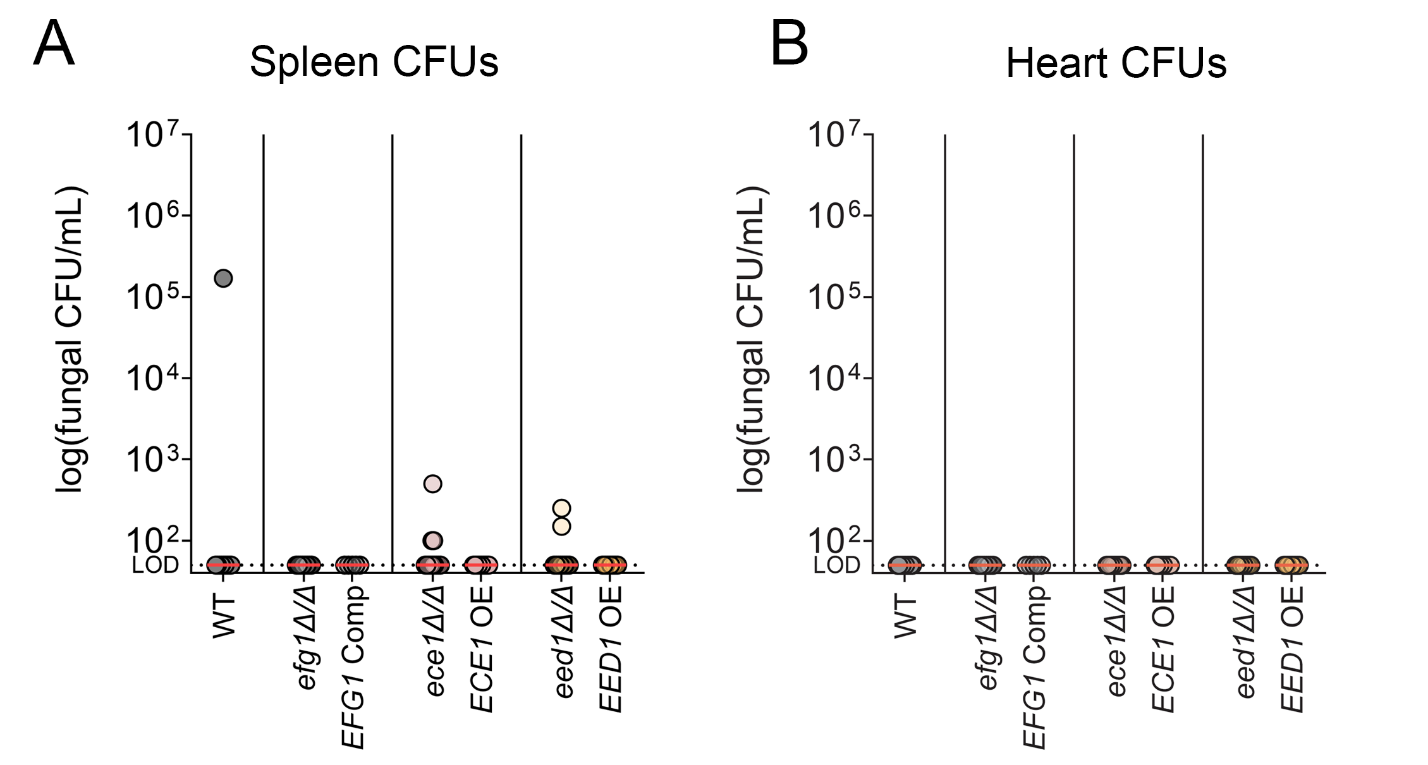

**Fig. S5. Spleen and heart colonization during mouse CAUTI.** SC5314 WT, SN425 *efg1Δ/Δ*, SN425 *EFG1* complement, SC5314 *ece1Δ/Δ*, SC5314 *ECE1* OE, SC5314 *eed1Δ/Δ*, and SC5314 *EED1* OE fungal burden in (**A**) spleen and (**B**) heart at 24 hours post infection and catheterization were quantitated as the number of CFUs recovered. For CFU enumeration, infections were done in at least two independent experiments with *n* = 3 or 4 mice for each one, and data are shown as the log(fungal CFU/organ or catheter). Animals that lost the catheter were not included in this work. The Mann-Whitney *U* test was used; *P* < 0.05 was considered statistically significant. **P* < 0.05, ***P* < 0.005, ****P* < 0.0005, and *****P* < 0.0001. The horizontal dotted line represents the limit of detection (LOD) of viable fungi. The horizontal red bar represents the median value.

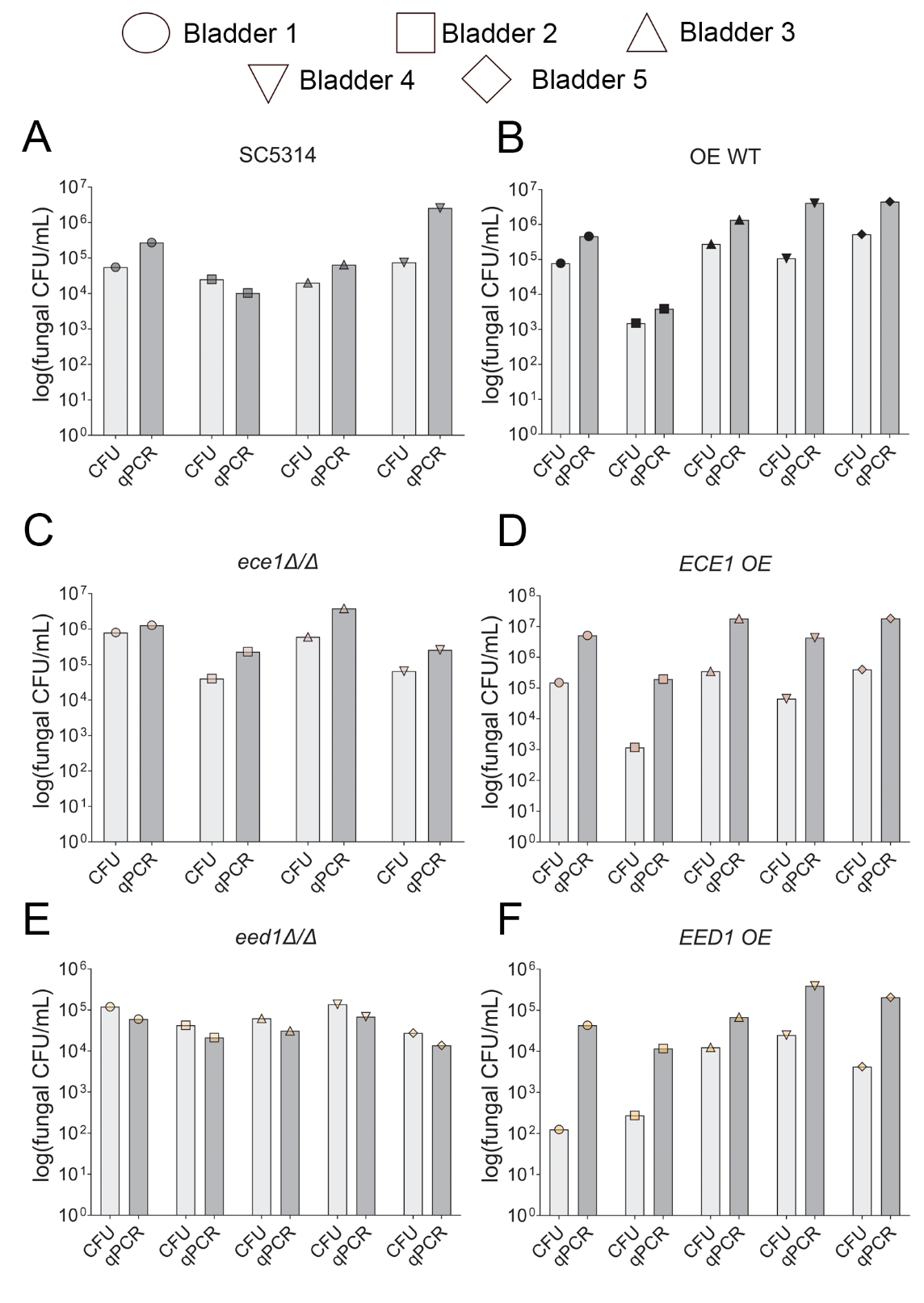

**Fig. S6. Bladder colonization during CAUTI quantified by qPCR compared to CFUs.** Mice were implanted and infected with ~1x10^6^ CFUs of strain (**A**) SC5314 WT, (**B**)OE WT, (**C**) SC5314 *ece1Δ/Δ*, (**D**) SC5314 *ECE1* OE, (**E**) SC5314 *eed1Δ/Δ*, or (**F**) SC5314 *EED1* OE. After 24h, harvested bladders were subjected to enumeration by colony forming units (CFU) and qPCR for comparison of fungal burden.

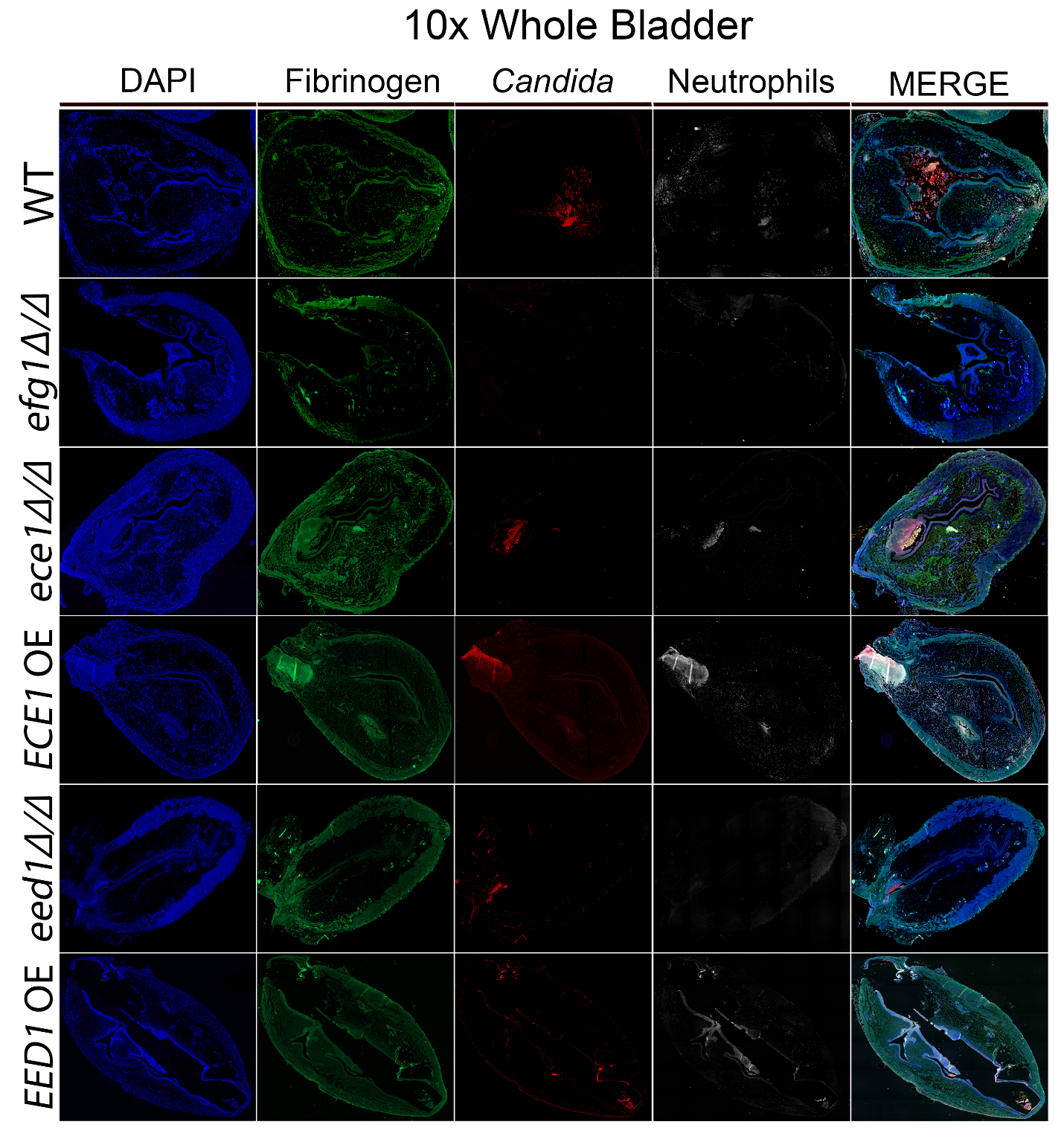

**Figure S7. Single channels of *C. albicans* WT and mutants bladder colonization during CAUTI at 10x magnification.** Mice were implanted and infected with ~1x 10^6^ CFUs with either strain SC5314 WT, SN250 *efg1Δ/Δ*, SC5314 *ece1Δ/Δ*, SC5314 *ECE1* OE, SC5314 *eed1Δ/Δ*, or SC5314 *EED1* OE At 24 hpi, bladder tissues were harvested, fixed, and parafilm-embedded. Bladders were subjected to IF analysis using antibodies to detect Fg (anti-Fg; green), *C. albicans* (anti-*Candida*; red), and neutrophils (anti-Ly6G; white). Staining with DAPI (blue) delineated the urothelium and cell nuclei (representative images). Magnification at 10x.

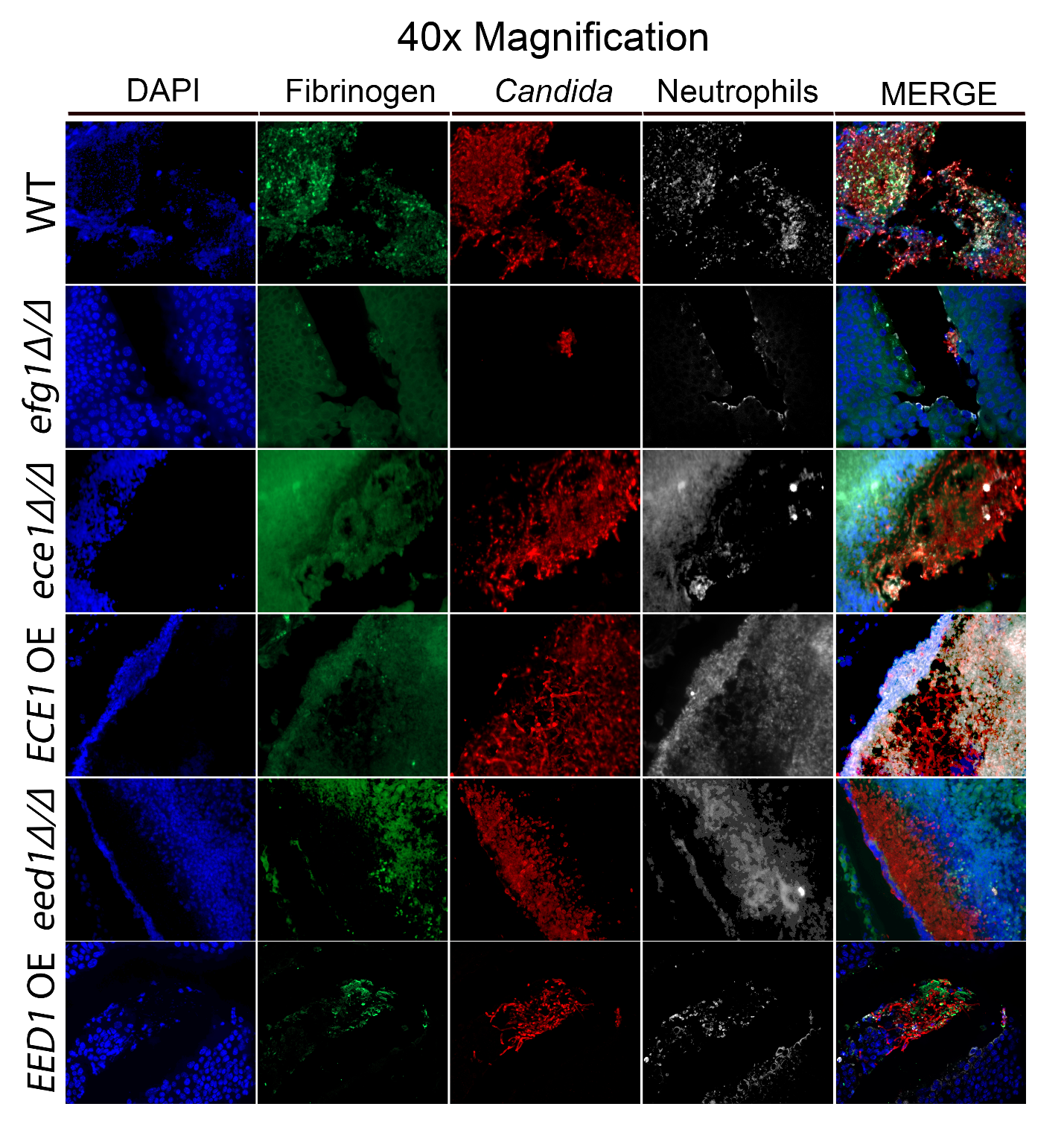

**Figure S8. Single channels of *C. albicans* WT and mutants bladder colonization during CAUTI at 40x magnification.** Mice were implanted and infected with ~1x10^6^ CFUs of strain SC5314 WT, SN250 *efg1Δ/Δ*, SC5314 *ece1Δ/Δ*, SC5314 *ECE1* OE, SC5314 *eed1Δ/Δ*, or SC5314 *EED1* OE. At 24 hpi, bladder tissues were harvested, fixed, and parafilm-embedded. Bladders were subjected to IF analysis using antibodies to detect Fg (anti-Fg; green), *C. albicans* (anti-*Candida*; red), and neutrophils (anti-Ly6G; white). Staining with DAPI (blue) delineated the urothelium and cell nuclei (representative images). Magnification at 40x.

**Table S1.** STRING network protein-protein node interactions of factors that promoted biofilm formation.

| **Node 1** | **Node 2** | **Neighborhood on chromosome** | **Gene fusion** | **Phylogenetic coocurrence** | **Homology** | **Coexpression** | **Experimentally determined interaction** | **Database annotated** | **Automated textmining** | **Combined score** |
| --- | --- | --- | --- | --- | --- | --- | --- | --- | --- | --- |
| CSA1 | RBT1 | 0 | 0 | 0 | 0 | 0 | 0 | 0 | 0.507 | 0.507 |
| HOG1 | SSK2 | 0 | 0 | 0.149 | 0.648 | 0.073 | 0.253 | 0 | 0.483 | 0.654 |
| HOG1 | NRG1 | 0 | 0 | 0 | 0 | 0 | 0.15 | 0.116 | 0.435 | 0.538 |
| NRG1 | SAP5 | 0 | 0 | 0 | 0 | 0 | 0.071 | 0 | 0.401 | 0.419 |
| NRG1 | RBT1 | 0 | 0 | 0 | 0 | 0 | 0.076 | 0.055 | 0.552 | 0.574 |

**Table S2.** STRING network protein-protein node interactions of factors with defective biofilm formation.

| **Node 1** | **Node 2** | **Neighborhood on chromosome** | **Gene fusion** | **Phylogenetic cooccurrence** | **Homology** | **Coexpression** | **Experimentally determined interaction** | **Database annotated** | **Automated textmining** | **Combined score** |
| --- | --- | --- | --- | --- | --- | --- | --- | --- | --- | --- |
| ADF1 | EFG1 | 0 | 0 | 0 | 0 | 0.336 | 0.12 | 0 | 0.328 | 0.573 |
| ADF1 | C1_07480C_A | 0 | 0 | 0 | 0 | 0 | 0 | 0 | 0.693 | 0.693 |
| ADR1 | PEX8 | 0 | 0 | 0 | 0 | 0 | 0.071 | 0 | 0.476 | 0.492 |
| ALI1 | NDU1 | 0 | 0 | 0 | 0 | 0.085 | 0.105 | 0.61 | 0.327 | 0.756 |
| ALI1 | C3_05550C_A | 0 | 0 | 0 | 0 | 0.112 | 0.426 | 0 | 0 | 0.468 |
| ALI1 | KIS1 | 0 | 0 | 0 | 0 | 0 | 0 | 0 | 0.617 | 0.617 |
| ALI1 | NUO2 | 0 | 0 | 0 | 0 | 0.38 | 0.752 | 0 | 0.34 | 0.889 |
| ALI1 | NUO1 | 0 | 0 | 0 | 0 | 0.239 | 0.82 | 0.54 | 0 | 0.931 |
| ALI1 | CR_02620C_A | 0 | 0 | 0 | 0 | 0.97 | 0.989 | 0.818 | 0.883 | 0.999 |
| ALI1 | C1_00700W_A | 0 | 0 | 0 | 0 | 0.819 | 0.989 | 0.818 | 0.534 | 0.999 |
| ALI1 | C6_02740W_A | 0 | 0 | 0 | 0 | 0.972 | 0.989 | 0.818 | 0.82 | 0.999 |
| ALI1 | C2_08650W_A | 0 | 0 | 0 | 0 | 0.797 | 0.989 | 0.647 | 0.63 | 0.999 |
| ALI1 | CR_10140W_A | 0.173 | 0 | 0 | 0 | 0.991 | 0.989 | 0.616 | 0.838 | 0.999 |
| ALI1 | MCI4 | 0 | 0 | 0 | 0 | 0.826 | 0.939 | 0.801 | 0.723 | 0.999 |
| ALS1 | ROB1 | 0 | 0 | 0 | 0 | 0.044 | 0.122 | 0 | 0.55 | 0.589 |
| ALS1 | CHT2 | 0 | 0 | 0 | 0 | 0 | 0 | 0 | 0.503 | 0.503 |
| ALS1 | PHR1 | 0 | 0 | 0 | 0 | 0 | 0.066 | 0 | 0.5 | 0.513 |
| ALS1 | CPH1 | 0 | 0 | 0 | 0 | 0 | 0.144 | 0.089 | 0.62 | 0.677 |
| ALS1 | BRG1 | 0 | 0 | 0 | 0 | 0.153 | 0.111 | 0 | 0.575 | 0.652 |
| ALS1 | PGA10 | 0 | 0 | 0 | 0 | 0 | 0 | 0 | 0.499 | 0.499 |
| ALS1 | EFG1 | 0 | 0 | 0 | 0 | 0 | 0.144 | 0.089 | 0.898 | 0.913 |
| ALS1 | LIP9 | 0 | 0 | 0 | 0 | 0 | 0 | 0 | 0.478 | 0.478 |
| APM1 | C2_04730W_A | 0 | 0 | 0 | 0 | 0 | 0 | 0 | 0.404 | 0.404 |
| APM1 | C3_02760C_A | 0 | 0 | 0 | 0 | 0 | 0.088 | 0.113 | 0.402 | 0.474 |
| APM1 | EVP1 | 0 | 0 | 0 | 0 | 0 | 0 | 0 | 0.402 | 0.402 |
| ARL1 | HET1 | 0 | 0 | 0 | 0 | 0.046 | 0.115 | 0.256 | 0.19 | 0.423 |
| AXL1 | POX18 | 0 | 0 | 0 | 0 | 0 | 0.073 | 0.598 | 0.065 | 0.621 |
| AXL1 | STE2 | 0 | 0 | 0 | 0 | 0.258 | 0.077 | 0 | 0.407 | 0.559 |
| AXL1 | PEX13 | 0 | 0 | 0 | 0 | 0.052 | 0 | 0.594 | 0 | 0.598 |
| BRG1 | ROB1 | 0 | 0 | 0 | 0 | 0.055 | 0.067 | 0 | 0.867 | 0.872 |
| BRG1 | YOX1 | 0 | 0 | 0 | 0 | 0 | 0.19 | 0.248 | 0.144 | 0.433 |
| BRG1 | CPH1 | 0 | 0 | 0 | 0 | 0.099 | 0.122 | 0.099 | 0.586 | 0.665 |
| BRG1 | CEK1 | 0 | 0 | 0 | 0 | 0.123 | 0.103 | 0 | 0.488 | 0.562 |
| BRG1 | EFG1 | 0 | 0 | 0 | 0 | 0 | 0.122 | 0.099 | 0.867 | 0.886 |
| BUR2 | SSN3 | 0 | 0 | 0 | 0 | 0 | 0.117 | 0.803 | 0.486 | 0.902 |
| CCN1 | PCL2 | 0 | 0 | 0 | 0 | 0.092 | 0.076 | 0 | 0.431 | 0.481 |
| CCN1 | NDU1 | 0 | 0 | 0 | 0 | 0.42 | 0 | 0 | 0 | 0.42 |
| CCN1 | GRR1 | 0 | 0 | 0 | 0 | 0 | 0.177 | 0 | 0.331 | 0.426 |
| CCN1 | CPH1 | 0 | 0 | 0 | 0 | 0 | 0.123 | 0 | 0.35 | 0.405 |
| CCN1 | CEK1 | 0 | 0 | 0 | 0 | 0 | 0.075 | 0.13 | 0.408 | 0.482 |
| CDC10 | CHS4 | 0 | 0 | 0 | 0 | 0 | 0.061 | 0 | 0.439 | 0.451 |
| CDC10 | GIN4 | 0 | 0 | 0 | 0 | 0.202 | 0.551 | 0 | 0.442 | 0.782 |
| CDG1 | FGR10 | 0 | 0 | 0 | 0 | 0 | 0 | 0 | 0.41 | 0.41 |
| CEK1 | FAV3 | 0 | 0 | 0 | 0 | 0.264 | 0 | 0 | 0.308 | 0.469 |
| CEK1 | MSB2 | 0 | 0 | 0 | 0 | 0.093 | 0.12 | 0 | 0.318 | 0.409 |
| CEK1 | DCK1 | 0 | 0 | 0 | 0 | 0 | 0 | 0 | 0.4 | 0.4 |
| CEK1 | FAR1 | 0 | 0 | 0 | 0 | 0.114 | 0.622 | 0.912 | 0.404 | 0.98 |
| CEK1 | STE2 | 0 | 0 | 0 | 0 | 0.307 | 0 | 0 | 0.648 | 0.746 |
| CEK1 | PHR1 | 0 | 0 | 0 | 0 | 0.124 | 0.122 | 0 | 0.401 | 0.499 |
| CEK1 | CPH1 | 0 | 0 | 0 | 0 | 0.325 | 0.763 | 0.907 | 0.884 | 0.998 |
| CEK1 | EFG1 | 0 | 0 | 0 | 0 | 0 | 0.147 | 0.116 | 0.696 | 0.751 |
| CEK1 | CHS7 | 0 | 0 | 0 | 0 | 0.094 | 0.183 | 0.239 | 0.244 | 0.517 |
| CEK1 | HST7 | 0 | 0 | 0.211 | 0.603 | 0.044 | 0.878 | 0.955 | 0.549 | 0.997 |
| CHS4 | YEA4 | 0 | 0 | 0 | 0 | 0 | 0 | 0 | 0.41 | 0.41 |
| CHS4 | GIN4 | 0 | 0 | 0 | 0 | 0.049 | 0.146 | 0 | 0.41 | 0.479 |
| CHS4 | CHS7 | 0 | 0 | 0 | 0 | 0.057 | 0 | 0 | 0.757 | 0.762 |
| CHS4 | CHS5 | 0 | 0 | 0 | 0 | 0 | 0 | 0 | 0.848 | 0.848 |
| CHS5 | YEA4 | 0 | 0 | 0 | 0 | 0 | 0 | 0 | 0.409 | 0.408 |
| CHS5 | CHS7 | 0 | 0 | 0 | 0 | 0.042 | 0 | 0 | 0.888 | 0.888 |
| CHS7 | STE2 | 0 | 0 | 0 | 0 | 0.044 | 0.312 | 0 | 0.182 | 0.414 |
| CHT2 | TOS1 | 0 | 0 | 0 | 0 | 0.084 | 0 | 0 | 0.418 | 0.444 |
| CHT2 | NGS1 | 0 | 0 | 0 | 0 | 0 | 0 | 0.9 | 0.316 | 0.928 |
| CHT2 | EFG1 | 0 | 0 | 0 | 0 | 0 | 0.065 | 0 | 0.403 | 0.417 |
| CHT2 | PHR1 | 0 | 0 | 0 | 0 | 0.06 | 0 | 0 | 0.504 | 0.513 |
| CHT2 | PIR1 | 0 | 0 | 0 | 0 | 0.047 | 0 | 0 | 0.559 | 0.561 |
| CHT2 | ENG1 | 0 | 0.006 | 0 | 0 | 0.451 | 0 | 0 | 0.422 | 0.669 |
| CHT2 | PGA4 | 0 | 0 | 0 | 0 | 0.081 | 0 | 0 | 0.671 | 0.684 |
| CPH1 | FAR1 | 0 | 0 | 0 | 0 | 0.07 | 0.191 | 0.111 | 0.408 | 0.551 |
| CPH1 | STE2 | 0 | 0 | 0 | 0 | 0.149 | 0.146 | 0 | 0.706 | 0.768 |
| CPH1 | EFG1 | 0 | 0 | 0 | 0 | 0 | 0.122 | 0 | 0.893 | 0.902 |
| CPH1 | HST7 | 0 | 0 | 0 | 0 | 0 | 0.191 | 0.113 | 0.906 | 0.926 |
| C6_00260W_A | ESC4 | 0 | 0 | 0 | 0 | 0 | 0.239 | 0.267 | 0.187 | 0.506 |
| C4_05580C_A | POX18 | 0 | 0 | 0 | 0 | 0.429 | 0.046 | 0 | 0 | 0.431 |
| C4_03600C_A | C5_03260C_A | 0 | 0 | 0 | 0 | 0 | 0.243 | 0.557 | 0.275 | 0.735 |
| C4_03600C_A | FAR1 | 0 | 0 | 0 | 0 | 0.051 | 0.243 | 0.557 | 0.275 | 0.738 |
| C4_03600C_A | PTC2 | 0 | 0 | 0 | 0 | 0 | 0.198 | 0.308 | 0.113 | 0.464 |
| C4_04360W_A | EHD3 | 0 | 0 | 0 | 0 | 0 | 0 | 0 | 0.628 | 0.628 |
| C4_04360W_A | CR_08920W_A | 0 | 0 | 0 | 0 | 0 | 0 | 0 | 0.628 | 0.628 |
| C2_04730W_A | STT4 | 0 | 0 | 0 | 0 | 0.043 | 0 | 0 | 0.405 | 0.406 |
| C2_04730W_A | C3_02760C_A | 0 | 0 | 0 | 0 | 0.1 | 0 | 0 | 0.402 | 0.438 |
| C2_08920W_ | TSC11 | 0 | 0 | 0 | 0 | 0 | 0.449 | 0 | 0.62 | 0.781 |
| CR_03430W_A | RCY1 | 0 | 0 | 0 | 0 | 0 | 0 | 0 | 0.613 | 0.613 |
| CR_03430W_A | STT4 | 0 | 0 | 0 | 0 | 0 | 0 | 0 | 0.479 | 0.479 |
| NDU1 | MCI4 | 0 | 0 | 0 | 0 | 0.08 | 0 | 0.56 | 0.088 | 0.598 |
| NDU1 | C6_02740W_A | 0 | 0 | 0 | 0 | 0.046 | 0 | 0.61 | 0.044 | 0.613 |
| NDU1 | C1_00700W_A | 0 | 0 | 0 | 0 | 0 | 0 | 0.61 | 0.115 | 0.64 |
| NDU1 | CR_02620C_A | 0 | 0 | 0 | 0 | 0.068 | 0 | 0.61 | 0.132 | 0.657 |
| C1_07480C_A | NPR2 | 0 | 0 | 0 | 0 | 0.055 | 0 | 0 | 0.596 | 0.602 |
| CR_02620C_A | C6_02740W_A | 0 | 0 | 0 | 0 | 0.989 | 0.989 | 0.986 | 0.889 | 0.999 |
| CR_02620C_A | PEX13 | 0 | 0 | 0 | 0 | 0 | 0 | 0 | 0.476 | 0.476 |
| CR_02620C_A | NUO1 | 0 | 0 | 0 | 0 | 0.703 | 0.82 | 0.54 | 0 | 0.973 |
| CR_02620C_A | CR_10140W_A | 0 | 0 | 0 | 0 | 0.761 | 0.989 | 0.266 | 0.826 | 0.999 |
| CR_02620C_A | KIS1 | 0 | 0 | 0 | 0 | 0.046 | 0 | 0 | 0.46 | 0.462 |
| CR_02620C_A | C2_08650W_A | 0 | 0 | 0 | 0 | 0.929 | 0.989 | 0.647 | 0.395 | 0.999 |
| CR_02620C_A | NUO2 | 0 | 0 | 0 | 0 | 0.689 | 0.648 | 0 | 0.474 | 0.937 |
| CR_02620C_A | MCI4 | 0 | 0 | 0 | 0 | 0.947 | 0.82 | 0.807 | 0.842 | 0.999 |
| CR_02620C_A | C1_00700W_A | 0 | 0 | 0 | 0 | 0.945 | 0.989 | 0.986 | 0.358 | 0.999 |
| C2_08650W_A | C6_02740W_A | 0 | 0 | 0 | 0 | 0.97 | 0.989 | 0.647 | 0.322 | 0.999 |
| C2_08650W_A | NUO1 | 0 | 0 | 0 | 0 | 0.21 | 0.82 | 0.54 | 0.294 | 0.947 |
| C2_08650W_A | CR_10140W_A | 0 | 0 | 0 | 0 | 0.688 | 0.989 | 0.598 | 0.618 | 0.999 |
| C2_08650W_A | NUO2 | 0 | 0 | 0 | 0 | 0.409 | 0.752 | 0 | 0.221 | 0.876 |
| C2_08650W_A | MCI4 | 0 | 0 | 0 | 0 | 0.879 | 0.82 | 0.642 | 0.343 | 0.994 |
| C2_08650W_A | C1_00700W_A | 0 | 0 | 0 | 0 | 0.937 | 0.989 | 0.647 | 0.536 | 0.999 |
| C6_02740W_A | NUO2 | 0 | 0 | 0 | 0 | 0.361 | 0.648 | 0 | 0.675 | 0.92 |
| C6_02740W_A | NUO1 | 0 | 0 | 0 | 0 | 0.13 | 0.82 | 0.54 | 0 | 0.921 |
| C6_02740W_A | MCI4 | 0 | 0 | 0 | 0 | 0.867 | 0.82 | 0.807 | 0.85 | 0.999 |
| C6_02740W_A | CR_10140W_A | 0 | 0 | 0 | 0 | 0.636 | 0.989 | 0.266 | 0.802 | 0.999 |
| C6_02740W_A | C1_00700W_A | 0 | 0 | 0 | 0 | 0.982 | 0.989 | 0.986 | 0.397 | 0.999 |
| C1_00700W_A | NUO1 | 0 | 0 | 0 | 0 | 0.828 | 0.82 | 0.54 | 0 | 0.984 |
| C1_00700W_A | CR_10140W_A | 0 | 0 | 0 | 0 | 0.493 | 0.989 | 0.266 | 0.69 | 0.998 |
| C1_00700W_A | MCI4 | 0 | 0 | 0 | 0 | 0.87 | 0.82 | 0.807 | 0.33 | 0.996 |
| C1_00700W_A | NUO2 | 0 | 0 | 0 | 0 | 0.28 | 0.648 | 0 | 0 | 0.736 |
| C7_00240W_A | C3_06150W_A | 0 | 0 | 0 | 0 | 0.045 | 0 | 0 | 0.691 | 0.692 |
| CR_08920W_A | EHD3 | 0 | 0 | 0 | 0 | 0.571 | 0.066 | 0 | 0.628 | 0.837 |
| UBC7 | C3_05680W_A | 0 | 0 | 0 | 0.776 | 0.044 | 0.602 | 0.96 | 0.266 | 0.987 |
| CR_00370W_A | GRR1 | 0 | 0 | 0 | 0 | 0.043 | 0.784 | 0.594 | 0.283 | 0.931 |
| CR_10140W_A | NUO1 | 0 | 0 | 0 | 0 | 0.521 | 0.82 | 0 | 0 | 0.91 |
| CR_10140W_A | EHD3 | 0 | 0 | 0 | 0 | 0.126 | 0 | 0 | 0.353 | 0.411 |
| CR_10140W_A | NUO2 | 0 | 0 | 0 | 0 | 0.679 | 0.752 | 0 | 0.592 | 0.964 |
| CR_10140W_A | MCI4 | 0 | 0 | 0 | 0 | 0.555 | 0.82 | 0.255 | 0.884 | 0.992 |
| DAC1 | OPT4 | 0 | 0 | 0 | 0 | 0.504 | 0 | 0 | 0 | 0.504 |
| DAC1 | NGS1 | 0 | 0 | 0 | 0 | 0 | 0 | 0 | 0.611 | 0.611 |
| EFG1 | ROB1 | 0 | 0 | 0 | 0 | 0.069 | 0.073 | 0 | 0.839 | 0.849 |
| EFG1 | PHR1 | 0 | 0 | 0 | 0 | 0 | 0.068 | 0 | 0.471 | 0.485 |
| EFG1 | HST7 | 0 | 0 | 0 | 0 | 0 | 0.121 | 0.113 | 0.576 | 0.64 |
| EHD3 | POX18 | 0.069 | 0 | 0 | 0 | 0.054 | 0.145 | 0.276 | 0.434 | 0.635 |
| EHD3 | PDK2 | 0 | 0 | 0 | 0 | 0.356 | 0.249 | 0 | 0.064 | 0.507 |
| EHD3 | PEX13 | 0 | 0 | 0 | 0 | 0.112 | 0 | 0.486 | 0.099 | 0.553 |
| ENG1 | PGA4 | 0 | 0 | 0 | 0 | 0.079 | 0 | 0 | 0.5 | 0.52 |
| FAR1 | HST7 | 0 | 0 | 0 | 0 | 0.044 | 0.248 | 0.159 | 0.617 | 0.737 |
| FAR1 | STE2 | 0 | 0 | 0 | 0 | 0.143 | 0.087 | 0 | 0.774 | 0.808 |
| GIN4 | TOS1 | 0 | 0 | 0 | 0 | 0.393 | 0.123 | 0 | 0 | 0.444 |
| GLN3 | YOX1 | 0 | 0 | 0 | 0 | 0 | 0.19 | 0.248 | 0.215 | 0.48 |
| GLN3 | STP2 | 0 | 0 | 0 | 0 | 0.044 | 0.121 | 0 | 0.492 | 0.535 |
| HST7 | STE2 | 0 | 0 | 0 | 0 | 0 | 0.077 | 0 | 0.841 | 0.847 |
| MCI4 | NUO1 | 0 | 0 | 0 | 0 | 0.377 | 0.866 | 0.54 | 0.432 | 0.975 |
| MCI4 | NUO2 | 0 | 0 | 0 | 0 | 0.857 | 0.752 | 0 | 0.695 | 0.988 |
| MNN14 | OCH1 | 0 | 0 | 0 | 0 | 0 | 0.072 | 0 | 0.501 | 0.517 |
| MSB2 | TOS1 | 0 | 0 | 0 | 0 | 0.334 | 0.148 | 0.081 | 0 | 0.432 |
| NUO1 | NUO2 | 0 | 0 | 0.21 | 0 | 0.153 | 0.648 | 0 | 0.88 | 0.967 |
| OCH1 | ZCF22 | 0 | 0 | 0 | 0 | 0 | 0.153 | 0.083 | 0.401 | 0.494 |
| PCL2 | YOX1 | 0 | 0 | 0 | 0 | 0.521 | 0 | 0 | 0.048 | 0.524 |
| PDK2 | RAD32 | 0.052 | 0 | 0 | 0 | 0.049 | 0 | 0 | 0.494 | 0.504 |
| PEX13 | POX18 | 0 | 0 | 0 | 0 | 0.224 | 0 | 0.971 | 0.324 | 0.983 |
| PGA10 | PHR1 | 0 | 0 | 0 | 0 | 0 | 0.066 | 0 | 0.403 | 0.418 |
| PGA4 | TOS1 | 0 | 0 | 0 | 0 | 0.116 | 0.066 | 0 | 0.496 | 0.548 |
| PGA4 | PIR1 | 0 | 0 | 0 | 0 | 0 | 0 | 0 | 0.676 | 0.676 |
| PHR1 | PIR1 | 0 | 0 | 0 | 0 | 0 | 0.044 | 0 | 0.399 | 0.4 |
| PIR1 | TOS1 | 0 | 0 | 0 | 0 | 0.049 | 0 | 0.049 | 0.674 | 0.679 |
| PTC2 | SSN3 | 0 | 0 | 0 | 0 | 0 | 0 | 0 | 0.475 | 0.475 |
| STT4 | TSC11 | 0 | 0 | 0 | 0 | 0 | 0 | 0 | 0.478 | 0.478 |

**Table S3.** Gene ontology slim analysis of significant biofilm formers (decreased biofilm formation, promoted biofilm formation).

| **GO-slim term** | **Deficient Biofilm** | **Promote Biofilm** | **Genes** |
| --- | --- | --- | --- |
| Transport | 64 | 6 | ALS1, APM1, ARL1, AXL1, C1_04150C_A, C1_07480C_A, C1_14200W_A, C1_05160C_A, C2_04750W_A, C2_07240C_A, C2_08920W_A, C2_09620W_A, C2_09860C_A, C3_05550C_A, C3_06150W_A, C4_01090C_A, C4_03050C_A, C4_03600C_A, C4_04360W_A, C7_00240W_A, CDC10, CDG1, CHS5, CHS7, CPH1, CR_01340W_A, CR_08920W_A, DAC1, DCK1, DCK2, DFG10, ECM14, ECM29, EHD3, EVP1, FAR1, FGR10, FPG1, GRR1, HAK1, HET1, IRF1, KAR3, KEX1, LIP4, LIP9, NPR2, OPT4, PDK2, PDR6, PEX13, PGA10, PGA49, PIR1, PLB3, RCY1, SSN3, STP2, STT4, TSC11, ULP2, YEA4, ZCF22, ZRC1  C1_12530C_A, C4_02920W_A, CSA1, HOG1, RBT1, SAP5 |
| Biological process regulation | 49 | 4 | AAF1, ADR1, ALS1, BRG1, BUR2, C2_00640W_A, C2_08920W_A, C3_02030W_A, C3_02760C_A, C3_03680W_A, C3_06150W_A, CAP1, CCN1, CEK1, CHS5, CHS7, CPH1, DCK1, DCK2, EFG1, ESC4, FAR1, GIN4, GLN3, GRR1, HST7, IRF1, KAR3, KIS1, MSB2, NGS1, NPR2, PCL2, PDK2, PEX8, PTC2, RAD32, ROB1, SAC7, SET3, SPT8, SSN3, STE2, STP2, TSC11, ULP2, YOX1, ZCF1, ZCF22  C7_02110W_A, HOG1, NRG1, SSK2 |
| Filamentous/hyphal growth | 37 | 3 | AAF1, ADR1, ALS1, APM1, ARL1, BRG1, C3_02760C_A, CCN1, CDC10, CEK1, CHS7, CHT2, CPH1, DAC1, DCK1, DCK2, DFG10, ECM29, EFG1, ESC4, FGR10, GIN4, GLN3, GRR1, HST7, KAR3, KIS1, MSB2, OCH1, PHR1, ROB1, RTK1, SET3, STP2, STT4, TSC11, ZRC1  HOG1, NRG1, SSK2 |
| Response to stress | 33 | 4 | ADR1, APM1, ARL1, BRG1, C3_02760C_A, C3_02770C_A, C3_03680W_A, CAP1, CEK1, CHT2, CPH1, DAC1, DCK1, DCK2, ECM29, EFG1, ESC4, FGR10, FPG1, GLN3, GRR1, HST7, KAR3, MSB2, OCH1, PRN1, RAD32, SET3, STP2, TSC11, UBC7, ULP2, ZRC1  RTN1, HOG1, NRG1, SSK2 |
| Response to chemical | 25 | 3 | ADR1, C2_01450C_A, C2_02520W_A, CAP1, CEK1, CPH1, CR_01340W_A, DAC1, EFG1, FAR1, GLN3, GRR1, HST7, MSB2, NGS1, NPR2, PHR1, PRN1, PTC2, SAC7, SSN3, STE2, STP2, UBC7, ZRC1  HOG1, NRG1, SSK2 |
| Interspecies interaction between organisms | 23 | 5 | AAF1, ALS1, BRG1, CAP1, CDC10, CEK1, CHS7, CHT2, CPH1, DAC1, EFG1, ENG1, GLN3, HET1, KEX1, LIP8, NGS1, NUO1, NUO2, PHR1, SET3, SSN3, ZCF1  ALS9, HOG1, NRG1, RBT1, SAP5 |
| Lipid metabolic process | 22 | 1 | ADR1, AXL1, C1_04150C_A, C2_07240C_A, C3_03680W_A, CCN1, CHS7, DFG10, FAD2, FAD3, HET1, LIP4, LIP8, LIP9, OCH1, PEX13, PGA10, PLB3, RCY1, STT4, TSC11, YOX1  RTN1 |
| Organelle organization | 21 | 1 | ADR1, ARL1, AXL1, C2_08920W_A, C3_02760C_A, C7_00240W_A, CDC10, DCK1, DCK2, GIN4, KAR3, LIP8, NDU1, NUO1, NUO2, PCL2, PEX13, RAD32, RTK1, STT4, TSC11  RTN1 |
| RNA metabolic process | 17 | 2 | BUR2, C1_00700W_A, C3_06150W_A, CHS4, CHS5, CR_02620C_A, FAV3, GIN4, KIS1, NGS1, PHR1, PTC2, RAD32, ROB1, SPT8, SSN3, ZRC1  C7_02110W_A, HOG1 |
| Unknown biological process | 17 | 1 | C1_04800C_A, C1_06250W_A, C2_00650W_A, C2_03490C_A, C2_04730W_A, C2_08650W_A, C2_09570C_A, C3_01180C_A, C4_05580C_A, C5_03260C_A, C6_00940C_A, C6_03310W_A, CIS301, IFA14, KRE62, POX18, TOS1  IFF6 |
| Cell wall organization | 14 | 2 | CDC10, CEK1, CHS5, CPH1, DCK1, ENG1, EVP1, HST7, MSB2, PGA4, PHR1, PIR1, TSC11, UBC7  HOG1, RBT1 |
| Vesicle-mediated transport | 14 | 3 | APM1, ARL1, C1_14200W_A, C4_03600C_A, C7_00240W_A, CHS5, CHS7, ECM29, GRR1, NPR2, PDR6, PGA10, RCY1, STT4  C1_12530C_A, C4_02920W_A, HOG1 |
| Cell cycle | 13 | 1 | AXL1, CCN1, CDC10, CHS5, ESC4, GIN4, GRR1, KAR3, NUO1, NUO2, PGA4, RAD32, ULP2  C7_02110W_A |
| Ribosome biogenesis | 13 | 0 | BUR2, C1_00700W_A, C3_06150W_A, CHS4, CR_02620C_A, FAV3, GIN4, NGS1, PEX13, PHR1, PTC2, RAD32, ZRC1 |
| Signal transduction | 13 | 2 | C3_03680W_A, CEK1, CPH1, DCK1, DCK2, FAR1, GIN4, HST7, KIS1, MSB2, SAC7, STE2, TSC11  HOG1, SSK2 |
| Biofilm formation | 10 | 4 | ALS1, BRG1, CCN1, CPH1, EFG1, GIN4, PGA10, PHR1, ROB1, STE2  ALS9, CSA1, NRG1, SAP5 |
| Protein catabolic process | 9 | 2 | C2_02520W_A, C3_02770C_A, C4_03050C_A, C4_03600C_A, CAP1, CR_00370W_A, ECM29, GRR1, UBC7  C4_02920W_A, SAP5 |
| Translation | 8 | 0 | ALI1, CHT2, CR_02620C_A, CR_03430W_A, FAD2, MCI4, NUO2, YOX1 |
| Cellular homeostasis | 8 | 2 | ALS1, C1_07480C_A, C2_00640W_A, CAP1, IRF1, OPT4, PGA10, ZRC1  CSA1, HOG1 |
| Cell adhesion | 7 | 2 | AAF1, ALS1, CHS7, EFG1, KIS1, PHR1, STE2  ALS9, NRG1 |
| Cytoskeleton organization | 6 | 0 | C2_08920W_A, CDC10, GIN4, KAR3, PCL2, TSC11 |
| Carbohydrate metabolic process | 5 | 0 | CHT2, CPH1, FAV3, NGS1, PGA4 |
| Pseudohyphal growth | 5 | 0 | CPH1, DFG10, EFG1, GRR1, HST7 |
| DNA metabolic process | 5 | 0 | C3_02770C_A, ESC4, FPG1, KAR3, RAD32 |
| Cytokinesis | 4 | 0 | AXL1, CDC10, CHS5, GIN4 |
| Generation of precursor metabolites and energy | 4 | 0 | C2_07240C_A, C6_02740W_A, CR_02620C_A, MCI4 |
| Cell development | 4 | 2 | CDC10, CHS5, EFG1, PGA4  HOG1, NRG1 |
| Cellular respiration | 3 | 0 | C6_02740W_A, CR_02620C_A, MCI4 |
| Protein folding | 2 | 0 | C3_05550C_A, CHS7 |
| Nucleus organization | 2 | 1 | C3_02760C_A, KAR3  RTN1 |
| Cell budding | 2 | 0 | AXL1, GIN4 |

**Table S4.** WT v *efg1∆/∆* in Fg-urine biofilms RNA sequencing hits with log2FoldChange greater than the absolute value of 2.

| **Gene** | **Log2FoldChange** | **-Log10pAdj** |
| --- | --- | --- |
| EFG1 | -7.18256 | 43.28773 |
| ARG4 | -6.85899 | 34.72112 |
| ECE1 | -5.1034 | 9.832439 |
| EED1 | -5.02956 | 30.4932 |
| C5_04980W_A | -4.83774 | 10.32569 |
| tR(UCU)4 | -4.57595 | 6.37387 |
| LEU2 | -4.51547 | 11.90791 |
| IFA14 | -4.31377 | 8.105058 |
| ALS1 | -4.11001 | 36.29284 |
| C3_04210W_A | -4.07934 | 5.328519 |
| C6_02450W_A | -4.03278 | 9.161389 |
| HGT6 | -4.02429 | 30.49121 |
| WH11 | -3.98058 | 5.562821 |
| C2_05180W_A | -3.82508 | 25.55081 |
| IHD1 | -3.82271 | 24.33447 |
| ATO1 | -3.63662 | 10.87346 |
| C3_00360W_A | -3.58172 | 29.65675 |
| FBA1 | -3.5785 | 43.38704 |
| BMT9 | -3.43077 | 6.990977 |
| SOD5 | -3.39956 | 1.708772 |
| C2_04000C_A | -3.39179 | 5.52996 |
| RHD1 | -3.37793 | 1.834369 |
| TEC1 | -3.37396 | 10.42811 |
| ZCF1 | -3.27768 | 7.530306 |
| RME1 | -3.24812 | 17.33793 |
| C3_02630C_A | -3.24076 | 6.890307 |
| WOR1 | -3.22915 | 3.867236 |
| WOR4 | -3.1343 | 13.46899 |
| CEK2 | -3.129 | 11.48169 |
| WOR3 | -3.12811 | 10.47546 |
| IRF1 | -3.08514 | 8.397692 |
| CTN3 | -3.05956 | 9.419213 |
| HGT8 | -3.04904 | 12.2104 |
| C6_00810C_A | -3.01855 | 9.677499 |
| HSP12 | -2.996 | 5.342254 |
| SSA2 | -2.98097 | 5.940285 |
| RBT4 | -2.97118 | 6.722865 |
| TYE7 | -2.95306 | 7.734199 |
| C1_13100W_A | -2.92734 | 7.106468 |
| C7_00880C_A | -2.86243 | 13.88447 |
| PGK1 | -2.85121 | 42.36913 |
| ADH5 | -2.7837 | 9.446437 |
| NAT4 | -2.75589 | 23.88043 |
| SLP3 | -2.74588 | 14.73373 |
| SOU2 | -2.73055 | 2.45681 |
| C3_03980C_A | -2.71458 | 9.147627 |
| C4_06620C_A | -2.7045 | 4.996822 |
| HSP21 | -2.69727 | 7.720642 |
| C7_03560W_A | -2.68343 | 0.898153 |
| CRD2 | -2.66697 | 5.352971 |
| HAK1 | -2.65831 | 3.926755 |
| HSP78 | -2.64515 | 9.678404 |
| PDC11 | -2.63957 | 19.16329 |
| DPP3 | -2.61639 | 4.813847 |
| C1_01620C_A | -2.60862 | 5.199559 |
| C2_08340C_A | -2.57449 | 5.171611 |
| C1_05150C_A | -2.57077 | 2.11269 |
| ACS1 | -2.52613 | 5.745033 |
| C2_07630C_A | -2.52506 | 13.33326 |
| ABP2 | -2.50878 | 2.761477 |
| HIS1 | -2.4933 | 4.803859 |
| C1_01630W_A | -2.4716 | 12.59369 |
| C2_00180C_A | -2.44292 | 3.988847 |
| C2_04670W_A | -2.41121 | 7.050341 |
| RNR22 | -2.40111 | 10.70144 |
| C1_04590W_A | -2.38347 | 5.397743 |
| tP(UGG)5 | -2.37957 | 1.488116 |
| C7_00350C_A | -2.35381 | 4.123579 |
| HAL9 | -2.34799 | 36.73494 |
| FRP3 | -2.33046 | 4.991197 |
| C1_02800W_A | -2.32242 | 4.898037 |
| C2_05770W_A | -2.30494 | 9.953457 |
| PHHB | -2.24797 | 9.939662 |
| C3_06860C_A | -2.24666 | 6.269546 |
| ACE2 | -2.24305 | 3.3963 |
| PST1 | -2.23829 | 3.857249 |
| C2_00510W_A | -2.23778 | 4.649801 |
| CR_07140C_A | -2.23573 | 4.857286 |
| RSN1 | -2.22387 | 5.883337 |
| C4_02740W_A | -2.22226 | 1.879465 |
| AAF1 | -2.18937 | 5.657025 |
| MRV4 | -2.1888 | 4.794555 |
| HSP30 | -2.17286 | 5.161712 |
| C6_02100W_A | -2.13335 | 3.995667 |
| CR_08880C_A | -2.13198 | 2.751013 |
| CR_04680C_A | -2.13026 | 2.125436 |
| C1_11670W_A | -2.12381 | 6.534999 |
| C2_08260W_A | -2.12023 | 2.276749 |
| CR_08420W_A | -2.11543 | 5.026002 |
| FGR23 | -2.0988 | 9.152628 |
| C6_03240W_A | -2.08652 | 11.2663 |
| ARR3 | -2.0849 | 3.397458 |
| C1_03750W_A | -2.06762 | 9.17951 |
| C6_03600C_A | -2.06223 | 2.333064 |
| C6_02620C_A | -2.03817 | 3.09119 |
| HSP70 | -2.03223 | 4.296405 |
| C1_06430C_A | -2.01536 | 1.482893 |
| UGA6 | -2.00515 | 4.21533 |
| AMO2 | -2.00149 | 1.339242 |
| SMC2 | -2.00072 | 4.137086 |
| ASH1 | 2.007109 | 3.104877 |
| GCY1 | 2.011434 | 1.30294 |
| tV(AAC)6 | 2.026302 | 2.807395 |
| HIS7 | 2.037591 | 3.696914 |
| MCM6 | 2.04628 | 4.92339 |
| AOX2 | 2.055829 | 2.452631 |
| RDN18 | 2.063097 | 2.110446 |
| C2_09180W_A | 2.068865 | 4.181371 |
| MXR1 | 2.070746 | 2.376933 |
| C3_02850C_A | 2.080445 | 7.727689 |
| HOM3 | 2.109931 | 2.655157 |
| CR_06550C_A | 2.112749 | 2.02599 |
| BIO2 | 2.125311 | 2.563805 |
| LYS4 | 2.127715 | 5.511768 |
| PGA54 | 2.13165 | 4.098052 |
| HGT12 | 2.151092 | 4.361067 |
| LYS9 | 2.151449 | 5.335865 |
| CTF8 | 2.159127 | 2.67913 |
| snR5d | 2.166292 | 5.983904 |
| C4_06910W_A | 2.190526 | 3.676247 |
| snR5a | 2.215136 | 4.483159 |
| C4_00210W_A | 2.215884 | 3.613717 |
| SNR52 | 2.216497 | 5.718766 |
| CIS308 | 2.226449 | 2.098067 |
| C5_04940W_A | 2.234493 | 2.658803 |
| tL(UAA)3 | 2.238521 | 1.396956 |
| C5_05180W_A | 2.251259 | 2.51331 |
| C7_03140W_A | 2.257297 | 6.436215 |
| ARO4 | 2.2757 | 7.211457 |
| PIR1 | 2.279299 | 2.045037 |
| HIS4 | 2.297834 | 7.001254 |
| CR_07300W_A | 2.308265 | 4.018038 |
| C1_06870C_A | 2.310319 | 4.312199 |
| CDC46 | 2.335235 | 3.201656 |
| C4_02080W_A | 2.340816 | 3.468138 |
| TER1 | 2.369511 | 16.06575 |
| TRP4 | 2.3761 | 3.683954 |
| PHR2 | 2.382906 | 1.82795 |
| SER33 | 2.398488 | 9.605689 |
| tH(GUG)1 | 2.469048 | 2.256374 |
| MCM3 | 2.476353 | 6.258198 |
| RBT5 | 2.558794 | 1.169099 |
| C6_04190C_A | 2.597703 | 11.38979 |
| CAT8 | 2.609163 | 3.892542 |
| IFG3 | 2.610933 | 8.21141 |
| C7_01010W_A | 2.644926 | 2.486329 |
| tY(GUA)2 | 2.654258 | 2.954196 |
| BNA31 | 2.68166 | 5.587222 |
| YMX6 | 2.68606 | 5.584055 |
| MET14 | 2.698299 | 3.857249 |
| tI(UAU)1 | 2.706022 | 3.619967 |
| C5_03040W_A | 2.736946 | 2.959346 |
| C3_04730C_A | 2.765183 | 8.384802 |
| CRH11 | 2.77382 | 3.397769 |
| tP(UGG)2 | 2.777692 | 5.352971 |
| CR_02060W_A | 2.789438 | 5.176864 |
| C5_03030W_A | 2.800154 | NA |
| CHT2 | 2.870723 | 8.900129 |
| PGA30 | 2.930689 | 1.925069 |
| HIS5 | 2.946874 | 6.478723 |
| C2_07840W_A | 2.960468 | 3.258124 |
| PRN3 | 2.99042 | 3.814705 |
| tL(CAA)6 | 3.012314 | 4.544874 |
| DAG7 | 3.01255 | 10.42529 |
| tY(GUA)3 | 3.052379 | 2.681976 |
| HOM6 | 3.068588 | 6.18526 |
| GRE3 | 3.071171 | 4.419301 |
| C4_01800W_A | 3.075098 | 2.23112 |
| tR(ACG)2 | 3.105976 | 5.221707 |
| CR_06500C_A | 3.115205 | 4.351925 |
| tL(AAG)1 | 3.117455 | 5.366039 |
| JEN2 | 3.21751 | 3.551348 |
| C1_09300C_A | 3.23795 | 5.352971 |
| HGT13 | 3.255857 | 5.641018 |
| IDP1 | 3.284019 | 6.736316 |
| tF(GAA)5 | 3.313063 | 3.259893 |
| ALS7 | 3.323875 | 43.28773 |
| CRP1 | 3.348381 | 3.677937 |
| CR_08310C_A | 3.366361 | 3.043635 |
| OAC1 | 3.39542 | 7.468582 |
| LEU1 | 3.440629 | 7.407589 |
| tR(ACG)1 | 3.452999 | 7.384852 |
| MET15 | 3.461053 | 4.73426 |
| MET3 | 3.519058 | 5.677684 |
| TNA1 | 3.534706 | 11.59382 |
| C1_06340W_A | 3.569739 | 7.045985 |
| PGA34 | 3.592859 | 33.10239 |
| RDN58 | 3.595356 | 1.793365 |
| CWP419 | 3.599348 | 6.592383 |
| C3_03570C_A | 3.617315 | 2.196671 |
| C6_02560W_A | 3.795874 | 5.44925 |
| PGA38 | 3.818613 | 6.76419 |
| tL(CAA)3 | 3.823187 | 5.745033 |
| ATX1 | 4.0388 | 2.135494 |
| LYS22 | 4.083506 | 11.09775 |
| ILV5 | 4.130104 | 9.884938 |
| LEU4 | 4.139637 | 7.689893 |
| C2_01630W_A | 4.234371 | 4.975925 |
| SCW11 | 4.248546 | 10.05418 |
| HSP31 | 4.283729 | 10.93965 |
| HGT17 | 4.292177 | 5.167112 |
| CR_07740W_A | 4.32828 | 13.80907 |
| tN(GUU)2 | 4.354852 | 6.473251 |
| CHT3 | 4.695553 | 9.154151 |
| HGT9 | 4.841102 | 12.12931 |
| C1_05830W_A | 4.843322 | 7.238047 |
| RBE1 | 4.988285 | 21.64387 |
| RHD3 | 5.109153 | 11.46254 |
| CR_01220W_A | 5.120526 | 3.857249 |
| XDJ1 | 5.174744 | 37.41543 |
| tK(UUU)3 | 5.189091 | 13.04783 |
| HGT10 | 5.248862 | 17.02524 |
| tR(UCU)5 | 5.308487 | 3.96304 |
| C4_01340W_A | 5.711526 | 8.179013 |
| PGA16 | 5.915479 | 7.059 |
| LDG3 | 5.933688 | 2.316158 |
| C7_02280W_A | 5.947416 | 10.96757 |
| tQ(UUG)4 | 5.983724 | 2.937495 |
| FGR41 | 6.055741 | 10.0472 |
| PGA31 | 6.459965 | 17.13342 |
| CR_06990W_A | 6.63725 | 22.87437 |
| tF(GAA)2 | 6.645589 | 9.216035 |
| C7_02260W_A | 6.701841 | 8.468085 |

**Table S5.** WT v *EFG1* complement in Fg-urine biofilms RNA sequencing hits with log2FoldChange greater than the absolute value of 2.

| **Gene** | **Log2FoldChange** | **-Log10pAdj** |
| --- | --- | --- |
| ARG4 | -9.1218 | 15.90344 |
| CR_10320W_A | -7.00996 | 15.07174 |
| C2_09800C_A | -6.26252 | 4.341581 |
| C4_05580C_A | -6.14979 | 33.86281 |
| C1_05970W_A | -6.09345 | 3.835997 |
| OP4 | -6.0357 | 12.74521 |
| C2_02230C_A | -5.97395 | 16.42224 |
| LEU2 | -5.92886 | 11.25081 |
| C1_05890W_A | -5.92464 | 21.54178 |
| SCW4 | -5.71733 | 13.23993 |
| JEN1 | -5.17254 | 1.458893 |
| PRY1 | -5.15553 | 6.722379 |
| C6_02450W_A | -4.57036 | 70.90772 |
| C7_02920W_A | -4.49343 | 65.61681 |
| POL93 | -4.16457 | 103.9495 |
| MRV2 | -4.14068 | 43.53257 |
| C1_00270W_A | -4.03912 | 101.3436 |
| PGA17 | -3.91239 | 16.21883 |
| C1_06430C_A | -3.84904 | 4.019438 |
| AMO2 | -3.67473 | 88.74445 |
| PXP2 | -3.65669 | 24.09917 |
| OFI1 | -3.5868 | 16.5381 |
| ATO1 | -3.31045 | 14.15881 |
| SOD5 | -3.3072 | 1.652293 |
| AGA1 | -3.29754 | 38.62998 |
| CR_07820W_A | -3.28492 | 10.38682 |
| CR_04220C_A | -3.25292 | 11.64111 |
| C2_08260W_A | -3.24129 | 3.305225 |
| NGT1 | -3.13377 | 3.765871 |
| GTT1 | -3.05417 | 2.617023 |
| ARO10 | -3.01315 | 3.524496 |
| CAR1 | -2.99651 | 59.80814 |
| CTN3 | -2.91568 | 62.3938 |
| GPT1 | -2.85845 | 10.3321 |
| DAL7 | -2.84868 | 10.47368 |
| FRP3 | -2.84086 | 19.21708 |
| WOR2 | -2.80333 | 1.495726 |
| HIS1 | -2.79655 | 24.3934 |
| ALK6 | -2.77378 | 5.419844 |
| ALK8 | -2.7237 | 6.670531 |
| EFG1 | -2.70801 | 10.20813 |
| MRV4 | -2.68869 | 11.81416 |
| ACS1 | -2.63121 | 35.66364 |
| C1_01630W_A | -2.59995 | 21.13525 |
| C2_01540W_A | -2.58037 | 29.89996 |
| C2_06990W_A | -2.53901 | 29.25821 |
| POX18 | -2.52244 | 8.347502 |
| CIT1 | -2.45188 | 2.405118 |
| C6_03240W_A | -2.44654 | 14.33858 |
| C5_03730W_A | -2.44424 | 6.754423 |
| PGA32 | -2.41248 | 8.242917 |
| CR_08920W_A | -2.40067 | 17.93655 |
| C6_02100W_A | -2.37609 | 5.401224 |
| C4_07000W_A | -2.36018 | 2.252367 |
| ATO10 | -2.35736 | 3.605166 |
| POX1-3 | -2.32292 | 42.80595 |
| CTR1 | -2.26119 | 10.7588 |
| C1_00830W_A | -2.25825 | 21.87504 |
| C7_00880C_A | -2.23972 | 18.36096 |
| C7_00870W_A | -2.21291 | 7.225761 |
| BLP1 | -2.20985 | 18.91143 |
| tR(UCU)4 | -2.20574 | 1.552134 |
| PGA45 | -2.19311 | 7.932004 |
| C1_02270C_A | -2.17609 | 12.57979 |
| C4_02190C_A | -2.16155 | 3.094416 |
| FAV1 | -2.15563 | 3.194301 |
| C4_07010C_A | -2.13983 | 17.44813 |
| GBU1 | -2.13686 | 7.766409 |
| FGR22 | -2.12348 | 3.642199 |
| C4_02930W_A | -2.09302 | 23.99528 |
| GTT13 | -2.09112 | 9.509387 |
| PCL1 | -2.06805 | 3.004771 |
| C4_03710C_A | -2.04256 | 9.756431 |
| C2_01450C_A | -2.00667 | 6.705467 |
| C2_00180C_A | -2.0046 | 1.797776 |
| ACU1 | 2.004126 | 11.72576 |
| PRN3 | 2.007455 | 21.89115 |
| TER1 | 2.015556 | 20.82099 |
| CRP1 | 2.022228 | 28.23222 |
| C7_01170C_A | 2.038026 | 20.18771 |
| GPX3 | 2.03819 | 5.888151 |
| MET14 | 2.0626 | 12.90195 |
| C1_09300C_A | 2.070951 | 9.759502 |
| TRP5 | 2.071887 | 29.42843 |
| LYS9 | 2.077872 | 13.28603 |
| FRE10 | 2.082696 | 9.314758 |
| PCL5 | 2.092799 | 12.72171 |
| CR_09010C_A | 2.100412 | 16.39038 |
| GIT1 | 2.112494 | 3.907533 |
| C3_01180C_A | 2.116644 | 17.75671 |
| FGR41 | 2.117139 | 3.747644 |
| C1_10360C_A | 2.119348 | 11.71135 |
| PGA31 | 2.126679 | 9.206706 |
| ACO2 | 2.129642 | 9.628755 |
| HIS7 | 2.137722 | 28.17031 |
| PHO114 | 2.138967 | 8.242917 |
| CHA1 | 2.139508 | 1.634845 |
| C4_06670W_A | 2.142609 | 16.20348 |
| MCM2 | 2.152369 | 12.30075 |
| ZCF25 | 2.155586 | 19.99963 |
| MCM3 | 2.169786 | 14.82407 |
| CDC46 | 2.176261 | 7.31706 |
| LEU42 | 2.203224 | 26.14615 |
| MXR1 | 2.208865 | 26.17685 |
| CDC54 | 2.231645 | 6.202051 |
| MRF1 | 2.234517 | 14.61881 |
| C4_04020C_A | 2.238646 | 3.210893 |
| BAT22 | 2.263251 | 19.4061 |
| RAS2 | 2.273091 | 13.26704 |
| MET6 | 2.291045 | 21.02078 |
| C1_08770W_A | 2.29577 | 29.2955 |
| OYE23 | 2.311764 | 9.756431 |
| CR_01220W_A | 2.417326 | 7.891679 |
| ILV3 | 2.417392 | 60.88545 |
| RAD10 | 2.426742 | 33.06174 |
| C1_06660W_A | 2.430451 | 25.61131 |
| AAT1 | 2.491636 | 14.18524 |
| HGT10 | 2.513246 | 13.31458 |
| ZWF1 | 2.530342 | 38.05583 |
| PGA13 | 2.539061 | 2.696076 |
| MCM6 | 2.55914 | 14.00192 |
| CR_06510W_A | 2.569925 | 15.97518 |
| SAP99 | 2.627615 | 14.25288 |
| ARO3 | 2.629658 | 27.62916 |
| TRP4 | 2.632706 | 17.21675 |
| YWP1 | 2.649646 | 29.56241 |
| GND1 | 2.670052 | 23.01598 |
| C1_01510W_A | 2.676939 | 9.780501 |
| HIS4 | 2.680792 | 52.01296 |
| CR_01630C_A | 2.692681 | 26.96996 |
| CR_02060W_A | 2.69324 | 24.45818 |
| QDR1 | 2.711668 | 11.67232 |
| BAT21 | 2.713328 | 35.80871 |
| YMX6 | 2.726693 | 18.92535 |
| MIS11 | 2.731377 | 48.54758 |
| MEP1 | 2.738748 | 27.72035 |
| CR_01440C_A | 2.739436 | 17.96386 |
| SNO1 | 2.761449 | 15.42904 |
| HIS5 | 2.80763 | 27.99227 |
| AOX2 | 2.843506 | 17.77551 |
| ILV2 | 2.858814 | 42.88224 |
| CAT8 | 2.865788 | 57.215 |
| SNZ1 | 2.899304 | 24.72355 |
| BIO2 | 2.96475 | 15.24282 |
| HOM3 | 2.982114 | 56.3271 |
| BNA31 | 3.002739 | 43.70055 |
| ATX1 | 3.006771 | 30.62638 |
| MET15 | 3.00807 | 47.97254 |
| MET3 | 3.054627 | 57.1347 |
| ILV6 | 3.08362 | 41.22296 |
| LYS4 | 3.084828 | 65.21175 |
| PHO87 | 3.129285 | 40.35009 |
| PHR2 | 3.161879 | 62.01571 |
| HOM6 | 3.350247 | 33.8163 |
| IFG3 | 3.413708 | 58.39738 |
| CR_08830W_A | 3.50959 | 3.355222 |
| C4_02080W_A | 3.697163 | 69.64815 |
| IDP1 | 3.778808 | 97.0646 |
| LYS22 | 3.783958 | 34.48143 |
| MAE1 | 3.980625 | 49.84827 |
| C1_12910W_A | 3.982768 | 64.61476 |
| OAC1 | 4.045935 | 58.96483 |
| CR_07740W_A | 4.123617 | 26.32775 |
| TNA1 | 4.33129 | 148.4137 |
| JEN2 | 4.356806 | 101.1606 |
| CR_06500C_A | 4.447401 | 35.55858 |
| ILV5 | 5.18958 | 117.9203 |
| LEU4 | 5.588561 | 181.911 |
| LEU1 | 5.610007 | 141.4946 |
| C4_01340W_A | 5.938641 | 26.03776 |
| C1_05830W_A | 6.428877 | 125.6201 |

**Table S6.** WT Fg-YPD v *EFG1* COMP Fg-YPD biofilms RNA sequencing hits with log2FoldChange greater than the absolute value of 2.

| **Gene** | **Log2FoldChange** | **-Log10pAdj** |
| --- | --- | --- |
| ARG4 | -9.606654992 | 11.9946864 |
| LEU2 | -7.382540258 | 21.20565818 |
| HIS1 | -4.72807365 | 37.93073598 |
| C2_04920W_A | -4.339932461 | 2.81291856 |
| C4_05580C_A | -4.275231599 | 4.368801796 |
| PXP2 | -4.27514 | 1.900563662 |
| JEN1 | -3.93242 | 3.03543373 |
| ATO1 | -3.750327753 | 3.565289585 |
| ACS1 | -3.23362 | 2.514954954 |
| C4_02230C_A | -3.09685 | 2.059640616 |
| C3_04310C_A | -2.89102 | 2.89495076 |
| SOU2 | -2.78827 | 1.168592014 |
| C4_04400C_A | -2.77661 | 0.978491417 |
| SCW4 | -2.762557801 | 6.61374213 |
| C6_01490C_A | -2.71492 | 1.702643524 |
| CAT1 | -2.577096292 | 2.484082573 |
| AGA1 | -2.54908 | 2.235509655 |
| FGR22 | -2.520864641 | 4.380251666 |
| CTN3 | -2.438406514 | 1.790771356 |
| FRP3 | -2.339748728 | 2.484082573 |
| CIT1 | -2.334652564 | 1.74618856 |
| C6_03600C_A | -2.309106548 | 1.881754841 |
| OP4 | -2.274990211 | 3.002120995 |
| HPD1 | -2.17801882 | 1.840493806 |
| ATO10 | -2.167545212 | 1.270729697 |
| ALK8 | -2.134624137 | 2.181497213 |
| CR_01840W_A | -2.121473773 | 0.642353387 |
| C2_02230C_A | -2.110747601 | 2.037807397 |
| ALD5 | -2.110406209 | 2.007404769 |
| CDC46 | 2.066718868 | 27.7303587 |
| ATX1 | 2.098495503 | 2.35879219 |
| C1_05830W_A | 2.963861492 | 5.490627521 |
| CR_08220C_A | 3.00157057 | 2.367156652 |

**Table S7.** WT Fg-YPD v *efg1∆/∆* Fg-YPD biofilms RNA sequencing hits with log2FoldChange greater than the absolute value of 2.

| **Gene** | **log2FoldChange** | **-log10padj** |
| --- | --- | --- |
| ARG4 | -9.02935 | 10.28988 |
| LEU2 | -8.24757 | 12.77211 |
| HIS1 | -4.80248 | 28.62342 |
| ATO1 | -3.89472 | 3.887864 |
| JEN1 | -3.74724 | 2.35819 |
| WOR1 | -3.67819 | 0.657434 |
| PXP2 | -3.25632 | 1.875922 |
| HSP12 | -3.1483 | 2.461221 |
| C6_01490C_A | -3.09634 | 2.191622 |
| C5_04980W_A | -3.07242 | 3.9694 |
| C6_03240W_A | -2.9708 | 7.320572 |
| C3_04310C_A | -2.80038 | 0.998334 |
| CAT1 | -2.64123 | 2.565202 |
| RME1 | -2.612 | 7.107349 |
| MAL31 | -2.51358 | 2.831151 |
| C1_05970W_A | -2.49597 | 1.132495 |
| ACS1 | -2.48399 | 1.454358 |
| INO1 | -2.36307 | 5.372634 |
| C6_03600C_A | -2.34531 | 2.875869 |
| FRP3 | -2.3309 | 2.483814 |
| C3_03570C_A | -2.32152 | 2.087708 |
| MAL2 | -2.29697 | 2.257188 |
| CTN3 | -2.21832 | 1.470115 |
| FAA21 | -2.17064 | 2.303006 |
| ALS1 | -2.15773 | 6.399027 |
| ASR1 | -2.14021 | 2.565202 |
| C2_01450C_A | -2.09286 | 2.354364 |
| SOU1 | -2.06322 | 2.305478 |
| C2_00180C_A | -2.01421 | 1.506999 |
| C1_02270C_A | -2.01135 | 1.845877 |
| C3_06040W_A | -2.01091 | 1.89742 |
| C4_00870W_A | 2.005812 | 1.04393 |
| C2_09180W_A | 2.006794 | 5.188425 |
| CR_03250C_A | 2.011913 | 2.346033 |
| PGA6 | 2.036258 | 6.627088 |
| C2_08580W_A | 2.045549 | 1.134663 |
| C7_01700W_A | 2.052317 | 1.470187 |
| CR_02320W_A | 2.057096 | 1.693011 |
| C7_03140W_A | 2.061226 | 1.086319 |
| CR_05410C_A | 2.061638 | 1.169196 |
| YOX1 | 2.061973 | 2.321409 |
| MNN4-4 | 2.068322 | #VALUE! |
| C1_00240C_A | 2.074488 | 1.112458 |
| SNR52 | 2.075741 | 1.946867 |
| C1_02340C_A | 2.088514 | 1.309047 |
| FLU1 | 2.094928 | 1.422065 |
| HSP104 | 2.100807 | 1.848173 |
| HCH1 | 2.113587 | 2.604358 |
| CR_02080W_A | 2.128848 | 1.21785 |
| C1_05580W_A | 2.157688 | 1.023833 |
| PGA53 | 2.190259 | 2.321409 |
| RNR1 | 2.198312 | 4.522879 |
| CR_05370W_A | 2.199529 | 1.845877 |
| C1_12500C_A | 2.208018 | 1.393398 |
| C5_00510W_A | 2.20979 | 9.183096 |
| C4_00900C_A | 2.233503 | 1.186678 |
| C2_04890C_A | 2.244303 | 1.129222 |
| ZRT1 | 2.245299 | 2.349301 |
| C1_00270W_A | 2.246552 | 1.410511 |
| C7_00350C_A | 2.248073 | 1.946867 |
| POL93 | 2.264132 | 1.410511 |
| C6_03740W_A | 2.28373 | 1.983172 |
| CR_07540C_A | 2.284032 | 2.028081 |
| WOR2 | 2.372265 | 2.360496 |
| SCW11 | 2.382792 | 1.182261 |
| C6_03370W_A | 2.391704 | 3.368496 |
| C7_00770W_A | 2.394541 | 2.215643 |
| CR_00030W_A | 2.396432 | 1.249777 |
| PTP2 | 2.401212 | 2.143906 |
| C1_05830W_A | 2.420241 | 2.557129 |
| CR_05070W_A | 2.430622 | 1.169196 |
| HSP31 | 2.478645 | 1.470115 |
| C1_06350W_A | 2.487284 | 2.078938 |
| OP4 | 2.529122 | 1.701713 |
| C3_05490C_A | 2.547261 | 1.520379 |
| CR_05850W_A | 2.551276 | 1.441663 |
| C1_10900C_A | 2.552669 | 1.627162 |
| FGR41 | 2.586874 | 2.000896 |
| C3_01240C_A | 2.589929 | 1.585455 |
| ACE2 | 2.617948 | 2.838855 |
| SWE1 | 2.62945 | 2.704826 |
| C5_04390C_A | 2.648048 | 1.222364 |
| PGA38 | 2.655163 | 2.008365 |
| FAV2 | 2.664959 | #VALUE! |
| ZRT2 | 2.697079 | 2.257188 |
| GIG1 | 2.700992 | 2.667301 |
| C4_05730W_A | 2.731475 | 1.842449 |
| PRA1 | 2.761248 | 2.565202 |
| C1_10050W_A | 2.772294 | 2.604358 |
| C1_07980C_A | 2.778741 | 2.312987 |
| PGA31 | 2.790488 | 3.50527 |
| CHT2 | 2.821063 | 2.305478 |
| C2_03710W_A | 2.833494 | 17.57675 |
| C6_04460W_A | 2.891234 | 1.750323 |
| SAP4 | 2.924557 | 1.458607 |
| C1_11710C_A | 2.933225 | 1.845877 |
| C3_02050C_A | 3.086293 | 1.296919 |
| CR_09630W_A | 3.111232 | 2.875869 |
| C1_06950C_A | 3.115045 | 2.453451 |
| CR_05600C_A | 3.194236 | 1.20164 |
| C2_10490C_A | 3.321659 | 2.604358 |
| RBR1 | 3.347889 | 1.013334 |
| NAG4 | 3.371724 | 2.124266 |
| HGT10 | 3.391244 | 3.148765 |
| C1_05310W_A | 3.46302 | 1.585455 |
| ALS7 | 3.505918 | 2.604358 |
| C2_01080C_A | 3.512146 | 2.161497 |
| C4_01110W_A | 3.635912 | 2.124266 |
| PGA34 | 3.698351 | 4.120904 |
| CR_08270W_A | 3.719127 | 3.527798 |
| C4_00550C_A | 3.864715 | 1.817109 |
| C3_04730C_A | 3.976123 | 3.552737 |
| C1_03570C_A | 4.069533 | 2.360496 |
| RBE1 | 4.254959 | 1.510381 |
| C4_00530C_A | 4.291512 | 2.565202 |
| C1_06340W_A | 4.338297 | 3.405182 |
| C1_11990W_A | 4.558556 | 3.749524 |
| CR_08220C_A | 4.755371 | 2.489672 |
| C1_03580W_A | 4.860134 | 3.787844 |
| CR_09390C_A | 5.83277 | 10.70115 |
| CR_06990W_A | 5.867012 | 1.931368 |

**Table S8.** Downregulated RNA sequencing hits in *efg1∆/∆* in urine and YPD when compared to WT.

| **Urine** | **YPD** |
| --- | --- |
| EFG1  ECE1  EED1  C5_04980W_A  IFA14  ALS1  C3_04210W_A  HGT6  WH11  C2_06570C_A  IHD1  C3_00360W_A  FBA1  BMT9  C2_04000C_A  RHD1  TEC1  ZCF1  RME1  C3_02630C_A  WOR1  WOR4  CEK2  WOR3  C2_08890W_A  HGT8  C6_00810C_A  HSP12  SSA2  RBT4  TYE7  C1_13100W_A  PGK1  ADH5  NAT4  CR_08990C_A  SOU2  C3_03980C_A  C4_06620C_A  HSP21  C7_03560W_A  CRD2  HAK1  HSP78  PDC11  DPP3  C1_01620C_A  C2_08340C_A  C1_05150C_A  C2_07630C_A  ABP2  C2_04670W_A  RNR22  C1_04590W_A  CR_05810C_A  C7_00350C_A  HAL9  C1_02800W_A  C2_05770W_A  PHHB  C3_06860C_A  ACE2  PST1  C2_00510W_A  CR_07140C_A  RSN1  C4_02740W_A  AAF1  HSP30  CR_08880C_A  CR_04680C_A  C1_11670W_A  CR_08420W_A  FGR23  ARR3  C1_03750W_A  C6_03600C_A  C6_02620C_A  HSP70  UGA6  C2_05860C_A | WOR1  HSP12  C5_04980W_A  C6_03240W_A  RME1  MAL31  C1_05970W_A  INO1  C3_03570C_A  MAL2  FAA21  ALS1  ASR1  C2_01450C_A  SOU1  C2_00180C_A  C1_02270C_A  C3_06040W_A |

**Table S8.** GO-slim ontology analysis of the Efg1-regulon in urine.

| **GO-slim Term** | **Observed gene count** | **Gene** |
| --- | --- | --- |
| Unknown biological process | 26 | C1_02800W_A, C1_04590W_A, C1_05150C_A, C1_11670W_A, C1_13100W_A, C2_00510W_A, C2_04000C_A, MSB2, C2_06570C_A, C2_08340C_A, C3_00360W_A, C3_02630C_A, C3_03980C_A, C3_04210W_A, C4_02740W_A, C4_06620C_A, C5_04980W_A, C6_00810C_A, C6_02620C_A, C7_00350C_A, CR_04680C_A, CR_07140C_A, CR_08880C_A, IFA14, IHD1, RSN1 |
| Transport | 25 | ADH5, ALS1, ARR3, BMT9, C1_01620C_A, C2_05770W_A, C6_03600C_A, CEK2, DPP3, ECE1, HAK1, HAL9, HGT6, HGT8, HSP21, HSP30, HSP70, IRF1, PHHB, PST1, SLP3, SOU2, SSA2, UGA6, WOR3 |
| Interspecies interaction between organisms | 19 | AAF1, ACE2, ALS1, EED1, ECE1, EFG1, FBA1, HSP21, HSP70, PGK1, PST1, RBT4, SSA2, TEC1, TYE7, WH11, WOR3, WOR4, ZCF1 |
| Biological process regulation | 16 | AAF1, ACE2, ALS1, C2_05770W_A, C2_05860C_A, EED1, EFG1, HAL9, IRF1, NAT4, RME1, TEC1, TYE7, WOR3, WOR4, ZCF1 |
| Filamentous/hyphal growth | 14 | AAF1, ACE2, ALS1, C1_03750W_A, C2_05770W_A, CEK2, EED1, EFG1, FGR23, HAL9, HSP21, PHHB, TEC1, TYE7 |
| Protein catabolic process | 13 | ARR3, C1_03750W_A, C2_07630C_A, C3_06860C_A, C5_02110W_A, CR_08420W_A, HSP12, RME1, RNR22, SLP3, SOU2, SSA2, WH11 |
| Response to stress | 13 | C3_06860C_A, C5_02110W_A, CEK2, EFG1, FGR23, HSP12, HSP21, HSP70, HSP78, PHHB, PST1, SLP3, TYE7 |
| Response to chemical | 11 | ACE2, C1_03750W_A, C3_06860C_A, C5_02110W_A, EFG1, HSP21, HSP70, PST1, SLP3, SSA2, TYE7 |
| Biofilm formation | 10 | ACE2, ADH5, ALS1, EED1, ECE1, EFG1, TEC1, TYE7, WH11, WOR3 |
| Cell adhesion | 8 | AAF1, ACE2, ALS1, C5_02110W_A, EED1, EFG1, TEC1, WOR4 |
| Carbohydrate metabolic process | 6 | C3_06860C_A, FBA1, HSP21, PDC11, PGK1, RHD1 |
| Cellular homeostasis | 5 | ALS1, ARR3, CRD2, IRF1, SOU2 |
| Lipid metabolic process | 4 | DPP3, FGR23, PDC11, RHD1 |
| Protein folding | 4 | CR_08420W_A, HSP70, HSP78, SSA2 |
| Generation of precursor metabolites and energy | 3 | FBA1, PDC11, PGK1 |
| Pseudohyphal growth | 2 | EFG1, TEC1 |
| Cell development | 2 | EFG1, PHHB |
| RNA metabolic process | 2 | C2_07630C_A, TYE7 |
| Translation | 2 | CRD2, tP(UGG)5 |
| Organelle organization | 2 | ABP2, HSP78 |
| Cytoskeleton organization | 1 | ABP2 |
| Cell wall organization | 1 | ACE2 |
| Cell cycle | 1 | PHHB |

**Table S9.** STRING network protein-protein node interactions of Efg1-regulon in urine.

| **Node1** | **Node2** | **Neighborhood on chromosome** | **Gene fusion** | **Phylogenetic cooccurrence** | **Homology** | **Co-**  **expression** | **Experimentally determined interaction** | **Database annotated** | **Automated textmining** | **Combined score** |
| --- | --- | --- | --- | --- | --- | --- | --- | --- | --- | --- |
| ACE2 | TYE7 | 0 | 0 | 0 | 0 | 0 | 0.119 | 0.064 | 0.41 | 0.47 |
| ACE2 | ALS1 | 0 | 0 | 0 | 0 | 0 | 0.144 | 0.089 | 0.48 | 0.559 |
| ACE2 | TEC1 | 0 | 0 | 0 | 0 | 0 | 0.171 | 0 | 0.554 | 0.614 |
| ACE2 | EFG1 | 0 | 0 | 0 | 0 | 0 | 0.071 | 0 | 0.624 | 0.636 |
| ADF1 | WOR3 | 0 | 0 | 0 | 0 | 0 | 0 | 0 | 0.469 | 0.469 |
| ADF1 | WOR4 | 0 | 0 | 0 | 0 | 0.044 | 0.12 | 0 | 0.506 | 0.548 |
| ADF1 | EFG1 | 0 | 0 | 0 | 0 | 0.336 | 0.12 | 0 | 0.328 | 0.573 |
| ADH5 | C1_11670W_A | 0 | 0 | 0 | 0 | 0.539 | 0 | 0 | 0 | 0.539 |
| ADH5 | C2_00510W_A | 0 | 0 | 0 | 0 | 0.47 | 0 | 0 | 0 | 0.47 |
| ADH5 | ECE1 | 0 | 0 | 0 | 0 | 0 | 0 | 0 | 0.403 | 0.403 |
| ADH5 | RSN1 | 0 | 0 | 0 | 0 | 0.415 | 0 | 0 | 0 | 0.414 |
| ADH5 | EFG1 | 0 | 0 | 0 | 0 | 0 | 0.092 | 0 | 0.404 | 0.435 |
| ADH5 | HGT6 | 0 | 0 | 0 | 0 | 0.475 | 0 | 0 | 0 | 0.475 |
| ADH5 | C3_06860C_A | 0.043 | 0 | 0 | 0 | 0.475 | 0 | 0 | 0.072 | 0.493 |
| ADH5 | C2_07630C_A | 0 | 0 | 0 | 0 | 0.454 | 0 | 0 | 0.221 | 0.556 |
| ADH5 | PDC11 | 0 | 0.048 | 0 | 0 | 0.054 | 0 | 0.9 | 0.1 | 0.908 |
| ALS1 | EED1 | 0 | 0 | 0 | 0 | 0.046 | 0.083 | 0.09 | 0.431 | 0.486 |
| ALS1 | ECE1 | 0 | 0 | 0 | 0 | 0 | 0 | 0 | 0.838 | 0.838 |
| ALS1 | EFG1 | 0 | 0 | 0 | 0 | 0 | 0.144 | 0.089 | 0.898 | 0.913 |
| ALS1 | IHD1 | 0 | 0 | 0 | 0 | 0.157 | 0.15 | 0.116 | 0.369 | 0.546 |
| ALS1 | TEC1 | 0 | 0 | 0 | 0 | 0.108 | 0.053 | 0 | 0.817 | 0.831 |
| CEK2 | EFG1 | 0 | 0 | 0 | 0 | 0 | 0.147 | 0.116 | 0.373 | 0.486 |
| CEK2 | TEC1 | 0 | 0 | 0 | 0 | 0 | 0.196 | 0 | 0.434 | 0.526 |
| C1_03750W_A | RNR22 | 0 | 0 | 0 | 0 | 0.537 | 0 | 0 | 0 | 0.537 |
| CR_07140C_A | C2_05770W_A | 0 | 0 | 0 | 0 | 0.046 | 0 | 0 | 0.68 | 0.681 |
| C1_11670W_A | HAK1 | 0 | 0 | 0 | 0 | 0 | 0 | 0 | 0.503 | 0.503 |
| C1_11670W_A | C5_04980W_A | 0 | 0 | 0 | 0 | 0.048 | 0 | 0 | 0.406 | 0.41 |
| C1_11670W_A 2 | PST1 | 0 | 0 | 0 | 0 | 0.488 | 0 | 0 | 0 | 0.487 |
| C1_11670W_A | RSN1 | 0 | 0 | 0 | 0 | 0.516 | 0 | 0 | 0 | 0.516 |
| C1_11670W_A | WH11 | 0 | 0 | 0 | 0 | 0.541 | 0 | 0 | 0 | 0.541 |
| C1_11670W_A | HSP12 | 0 | 0 | 0 | 0 | 0.677 | 0 | 0 | 0 | 0.677 |
| C1_11670W_A | C2_07630C_A | 0 | 0 | 0 | 0 | 0.755 | 0 | 0 | 0 | 0.755 |
| C2_07630C_A | C2_00510W_A | 0 | 0 | 0 | 0 | 0.612 | 0 | 0 | 0 | 0.612 |
| C2_07630C_A | WH11 | 0 | 0 | 0 | 0 | 0.678 | 0 | 0 | 0 | 0.678 |
| C2_07630C_A | PST1 | 0 | 0 | 0 | 0 | 0.477 | 0 | 0 | 0 | 0.477 |
| C2_07630C_A | HSP12 | 0 | 0 | 0 | 0 | 0.754 | 0 | 0 | 0 | 0.754 |
| C2_07630C_A | C7_00350C_A | 0 | 0 | 0 | 0 | 0.493 | 0 | 0 | 0 | 0.493 |
| C2_07630C_A | RSN1 | 0 | 0 | 0 | 0 | 0.553 | 0 | 0 | 0 | 0.553 |
| C2_07630C_A | C3_06860C_A | 0 | 0 | 0 | 0 | 0.663 | 0 | 0 | 0 | 0.663 |
| C2_00510W_A | RSN1 | 0 | 0 | 0 | 0 | 0.589 | 0 | 0 | 0 | 0.589 |
| C3_02630C_A | ZCF1 | 0 | 0.547 | 0 | 0 | 0.559 | 0 | 0 | 0 | 0.792 |
| C3_02630C_A | FGR23 | 0 | 0 | 0 | 0 | 0.046 | 0.368 | 0 | 0.097 | 0.408 |
| C1_01620C_A | HSP30 | 0 | 0 | 0 | 0 | 0.066 | 0.065 | 0 | 0.478 | 0.504 |
| CR_08420W_A | C7_00350C_A | 0 | 0 | 0 | 0 | 0.745 | 0 | 0 | 0.072 | 0.753 |
| CR_08420W_A | SSA1 | 0.043 | 0 | 0.312 | 0 | 0.394 | 0.196 | 0.253 | 0.092 | 0.732 |
| CR_08420W_A | SSA2 | 0.043 | 0 | 0.32 | 0 | 0.359 | 0.196 | 0.253 | 0.092 | 0.719 |
| CR_08420W_A | HSP78 | 0.045 | 0 | 0 | 0 | 0.526 | 0.148 | 0 | 0.117 | 0.614 |
| C3_06860C_A | RNR22 | 0 | 0 | 0 | 0 | 0.676 | 0 | 0 | 0 | 0.676 |
| C3_06860C_A | C7_00350C_A | 0 | 0 | 0 | 0 | 0.508 | 0.065 | 0 | 0 | 0.52 |
| C3_06860C_A | HGT6 | 0 | 0 | 0 | 0 | 0.473 | 0 | 0 | 0 | 0.473 |
| C3_06860C_A | RSN1 | 0 | 0 | 0 | 0 | 0.473 | 0 | 0 | 0 | 0.473 |
| C7_00350C_A | SSA1 | 0 | 0 | 0 | 0 | 0.093 | 0.115 | 0.263 | 0.135 | 0.42 |
| EED1 | TEC1 | 0 | 0 | 0 | 0 | 0.046 | 0 | 0 | 0.592 | 0.594 |
| EED1 | DPP3 | 0 | 0 | 0 | 0 | 0.044 | 0.124 | 0 | 0.591 | 0.627 |
| EED1 | ECE1 | 0 | 0 | 0 | 0 | 0.093 | 0 | 0 | 0.649 | 0.668 |
| EED1 | EFG1 | 0 | 0 | 0 | 0 | 0.126 | 0.148 | 0.06 | 0.673 | 0.74 |
| DPP3 | EFG1 | 0 | 0 | 0 | 0 | 0 | 0.119 | 0 | 0.397 | 0.446 |
| ECE1 | RBT4 | 0 | 0 | 0 | 0 | 0.057 | 0 | 0 | 0.612 | 0.618 |
| ECE1 | TEC1 | 0 | 0 | 0 | 0 | 0.045 | 0 | 0 | 0.693 | 0.694 |
| ECE1 | IHD1 | 0 | 0 | 0 | 0 | 0.28 | 0 | 0 | 0.618 | 0.713 |
| ECE1 | EFG1 | 0 | 0 | 0 | 0 | 0 | 0 | 0 | 0.863 | 0.863 |
| EFG1 | WH11 | 0 | 0 | 0 | 0 | 0.066 | 0 | 0 | 0.56 | 0.571 |
| EFG1 | RBT4 | 0 | 0 | 0 | 0 | 0.089 | 0.043 | 0 | 0.395 | 0.426 |
| EFG1 | WOR4 | 0 | 0 | 0 | 0 | 0.046 | 0 | 0 | 0.58 | 0.582 |
| EFG1 | TYE7 | 0 | 0 | 0 | 0 | 0.228 | 0.119 | 0.055 | 0.506 | 0.64 |
| EFG1 | WOR3 | 0 | 0 | 0 | 0 | 0 | 0.074 | 0 | 0.678 | 0.689 |
| EFG1 | TEC1 | 0 | 0 | 0 | 0 | 0 | 0.152 | 0 | 0.942 | 0.949 |
| FBA1 | SSA1 | 0 | 0 | 0 | 0 | 0.156 | 0 | 0 | 0.35 | 0.428 |
| FBA1 | PGK1 | 0.115 | 0 | 0 | 0 | 0.999 | 0.166 | 0 | 0.95 | 0.999 |
| FBA1 | PDC11 | 0 | 0 | 0 | 0 | 0.999 | 0 | 0.8 | 0.713 | 0.999 |
| HGT6 | RNR22 | 0 | 0 | 0 | 0 | 0.472 | 0 | 0 | 0 | 0.472 |
| HSP12 | WH11 | 0 | 0 | 0.057 | 0.901 | 0.999 | 0 | 0 | 0.068 | 0.999 |
| HSP12 | SSA2 | 0 | 0 | 0 | 0 | 0.05 | 0.119 | 0 | 0.475 | 0.522 |
| HSP12 | HSP30 | 0 | 0 | 0 | 0 | 0.06 | 0 | 0 | 0.513 | 0.523 |
| HSP12 | HSP78 | 0 | 0 | 0 | 0 | 0.082 | 0 | 0 | 0.652 | 0.667 |
| HSP12 | SSA1 | 0 | 0 | 0 | 0 | 0.109 | 0.119 | 0 | 0.644 | 0.696 |
| HSP12 | HSP21 | 0 | 0 | 0 | 0 | 0.091 | 0 | 0 | 0.713 | 0.728 |
| HSP21 | SSA1 | 0.055 | 0 | 0 | 0 | 0.484 | 0.079 | 0 | 0.7 | 0.847 |
| HSP21 | SSA2 | 0.055 | 0 | 0 | 0 | 0.322 | 0.079 | 0 | 0.535 | 0.688 |
| HSP21 | HSP78 | 0.095 | 0 | 0 | 0 | 0.729 | 0.149 | 0 | 0.4 | 0.858 |
| HSP30 | SSA1 | 0 | 0 | 0 | 0 | 0.044 | 0 | 0 | 0.598 | 0.599 |
| HSP30 | RME1 | 0 | 0 | 0 | 0 | 0.476 | 0 | 0 | 0 | 0.476 |
| HSP30 | HSP78 | 0 | 0 | 0 | 0 | 0.066 | 0 | 0 | 0.837 | 0.841 |
| HSP78 | SSA1 | 0.055 | 0 | 0 | 0 | 0.649 | 0.27 | 0 | 0.713 | 0.921 |
| HSP78 | SSA2 | 0.055 | 0 | 0 | 0 | 0.496 | 0.27 | 0 | 0.492 | 0.8 |
| IHD1 | WOR3 | 0 | 0 | 0 | 0 | 0.082 | 0.043 | 0.274 | 0.285 | 0.483 |
| NAT4 | PHHB | 0 | 0 | 0 | 0 | 0 | 0 | 0 | 0.403 | 0.403 |
| PDC11 | SSA1 | 0 | 0 | 0 | 0 | 0.095 | 0 | 0 | 0.479 | 0.508 |
| PDC11 | PGK1 | 0 | 0 | 0 | 0 | 0.891 | 0 | 0 | 0.768 | 0.973 |
| PDC11 | SSA2 | 0 | 0 | 0 | 0 | 0.044 | 0 | 0 | 0.409 | 0.41 |
| PGK1 | SSA1 | 0 | 0 | 0 | 0 | 0.125 | 0.117 | 0 | 0.605 | 0.668 |
| PGK1 | SSA2 | 0 | 0 | 0 | 0 | 0.071 | 0.117 | 0 | 0.555 | 0.603 |
| RHD1 | RME1 | 0 | 0 | 0 | 0 | 0.058 | 0 | 0 | 0.406 | 0.416 |
| RNR22 | SLP3 | 0 | 0 | 0 | 0 | 0.403 | 0 | 0 | 0 | 0.403 |
| RNR22 | RSN1 | 0 | 0 | 0 | 0 | 0.529 | 0 | 0 | 0 | 0.529 |
| SSA1 | SSA2 | 0 | 0 | 0.049 | 0.983 | 0.051 | 0.23 | 0.9 | 0.055 | 0.922 |
| TEC1 | TYE7 | 0 | 0 | 0 | 0 | 0 | 0 | 0 | 0.475 | 0.475 |
| WOR3 | WOR4 | 0 | 0 | 0 | 0 | 0 | 0.074 | 0 | 0.863 | 0.867 |

**Table S10.** Strains used in this study.

| **Species** | **Strain** | **Description** | **Reference** |
| --- | --- | --- | --- |
| *Candida albicans* | SC5314 | Laboratory strain | Odds *et al.* 2004 |
| *Candida albicans* | *efg1∆/∆* (SN250 background; CJN2302) | Laboratory strain from Clarissa J. Nobile | Nobile *et al.* 2012 |
| *Candida albicans* | *EFG1* complement (SN250 background; CJN2318) | Laboratory strain from Clarissa J. Nobile | Nobile *et al.* 2012 |
| *Candida albicans* | *efg1∆/∆* (SC5314 background) | Laboratory strain from Rebecca Shapiro. | This study |
| *Candida albicans* | *ece1∆/∆* (SC5314 background) | Laboratory strain from Rebecca Shapiro. | This study |
| *Candida albicans* | *eed1∆/∆* (SC5314 background) | Laboratory strain from Rebecca Shapiro. | This study |
| *Candida albicans* | *ECE1* Overexpression (SC5314 background) | Laboratory strain from Rebecca Shapiro. | This study |
| *Candida albicans* | *EED1* Overexpression (SC5314 background) | Laboratory strain from Rebecca Shapiro. | This study |
| *Candida albicans* | Overexpression Control | Laboratory strain from Rebecca Shapiro. |  |

**Table S11.** Primers used in this study.

| **Target** | **Forward primer (5’-3’)** | **Reverse primer (5’-3’)** |
| --- | --- | --- |
| *ITS2* | TGGGTTTGCTTGAAAGACGG | CCGCCGCAAGCAATGTTTTT |
| *EFG1* | GGTCAGTATAATGCTCCTGGTAAG | CAGCACCACCCTGGTAATAAT |
| *ECE1* | AGCTGACCAAGCACCTACTG | TCTGACGACGGCATTAGCAA |
| *EED1* | TCTCGTGGTTCCAATGCCAG | GGTGGTACTGGTCCACCTCT |
| *FBA1* | TGGCTCCTCCAGCAGTTTTA | ATGTAGTGAGCGGCAGCAAT |
| *HGT6* | ACTGGTTGTGGGGTTTCTTGA | CTTGCTCTTCATCTGCGTGG |
| *TYE7* | AGAACCAGGTACGAAGGCAG | CAAAAGCCACCCAAGGAATGA |
| *HSP21* | TCCGACAAATACGTGGTTTCCT | ACTTGGACACCTTGCTCACC |
| *PGK1* | TTCGATGAAGCCGGTGCTAA | ACCACAGTCCAAACCCATCC |
| *PDC11* | ATCTTGGTTGATGCTTGTGCC | ACCAACGTAAACACCACCGA |
| *C4_02740W_A* | TCCCACAACTCAACCTCCAC | ACATCAAAGCAGTAGTCGTGC |
| *ALS1* | GACTAGTGAACCAACAAATACCAG | ACCAGAAGAAACAGCAGGTG |
| *PGA34* | TAGTGCTGCTTCCCTTGCTT | CAGTTTTTGTAGTCACAGCAGCA |
| *ALS7* | TCCAAGTACTTCTGTACCATCGAG | CGAAAGTTCCCCCACGGTAA |
